# The latent cis-regulatory potential of mobile DNA

**DOI:** 10.1101/2023.10.22.563463

**Authors:** Timothy Fuqua, Andreas Wagner

**Affiliations:** Department of Evolutionary Biology and Environmental Studies, University of Zurich, Zurich, Switzerland; Swiss Institute of Bioinformatics, Quartier Sorge-Batiment Genopode, Lausanne, Switzerland; The Sante Fe Institute, Sante Fe, NM, USA

## Abstract

Transposable elements can alter gene regulation in their host genome, either when they integrate into a genome, or when they accrue mutations after integration. However, the extent to which transposons can alter gene expression, as well as the necessary mutational steps, are not well characterized. Here we study the gene regulatory potential of the prominent IS3 family of transposable elements in *E.coli*. We started with 10 sequences from the ends of 5 IS3 sequences, created 18,537 random mutations in them, and measured their promoter activity using a massively parallel reporter assay. All 10 sequences could evolve de-novo promoter activity from single point mutations. De-novo promoters mostly emerge from existing proto-promoter sequences when mutations create new −10 boxes downstream of preexisting −35 boxes. The ends of IS3s harbor ~1.5 times as many such proto-promoter sequences than the *E. coli* genome. We also estimate that at least 26% of the 706 characterized IS3s already encode promoters. Our study shows that transposable elements can have a high latent cis-regulatory potential. Our observations can help to explain why mobile DNA may persist in prokaryotic genomes. They also underline the potential use of transposable elements as a substrate for evolving new gene expression.

## Introduction

Insertion Sequences (ISs) are simple mobile genetic elements that encode a transposase flanked by inverted repeat sequences that serve as binding sites for the transposase^1^. There are various families of ISs, which differ in the number of open reading frames they encode, and the mechanisms by which they transpose^1,2^. One of the best characterized *E. coli* IS families is IS3, with over 700 characterized members. IS3s are ~1 kilobase (kb) in length and contain 2 ORFs (*orfA* and *orfB*) which regulate transposition^3^. The ORFs fuse via translational frameshifting to create the transposase (*orfAB*)^4,5^. Transposition works through a “copy-paste” mechanism using a DDE-transposase^1^.

Canonical prokaryotic promoters encode two motifs called the −35 and −10 boxes^6^, which are spaced 15-19 base pairs apart^7^, and serve as a binding site for the σ_70_ subunit of the RNA polymerases that transcribes genes^8^. Protein transcription factors (TFs) may bind near such promoter sequences to activate or repress transcription^9^. Additional σ subunits are expressed under different environmental conditions and bind to their own unique motifs^10^. In members of the IS3 family, a native promoter is usually located on the left end to transcribe the ORFs^3^. Additionally, after excision, IS3s form circular DNA intermediary “minicircles” that join the left and right ends^5^. When the right end encodes a −35 box and the left end a −10 box, a strong “junction promoter” can result from this joining^11,12^.

In addition to their native promoter, ISs can also contain partial or complete outward-directed promoters^13,14^. An IS with a complete outward-directed promoter can transcribe adjacent genes^15–17^. In addition, an IS can also serendipitously integrate upstream of a genomic −10 box, enabling it to form a “hybrid promoter” with its own −35 box^1,13,18–20^. Furthermore, de-novo promoter activity has been described in ISs upon mutation^21^. Outward-directed promoter activity in ISs has led to many evolutionary innovations^22,23^, including the evolution of antibiotic resistance^19,20^.

These observations raise the possibility that ISs have a high inherent gene regulatory potential. However, recent studies show that even random DNA sequences can have such a potential. For example, one study in *E. coli* showed that ~10% of randomly synthesized DNA encodes promoters^8,24–26^, and single mutations to non-promoter DNA can be sufficient to create new promoters^8,24,27^. It is thus not clear whether the regulatory potential of IS families like IS3 is especially high, because promoters in IS3s were not identified systematically, and are only described in few IS3s^2^.

To evaluate the regulatory potential of IS3s, we created over 18,000 random mutations to 10 ends of 5 IS3 sequences, and measured their promoter activity using a massively parallel reporter assay. De-novo promoters emerge from IS3s in or around preexisting −10 and −35 boxes, primarily when single mutations create new −10 boxes downstream of preexisting −35 boxes. We find that the ends of IS3s are enriched with such proto-promoter sequences, encoding ~1.5× as many compared to the *E. coli* genome. We also estimate that at least 26% of the 706 characterized IS3s already encode complete outward-directed promoters. Overall, our study shows that IS3s do indeed have a high latent cis-regulatory potential, which could help to explain why mobile DNA persists in prokaryotic genomes.

## Results

### Promoters emerge from the ends of IS3s at different probabilities

To choose an appropriate size of DNA fragments for mutagenesis, we first performed a preliminary experiment that inserted a functional promoter at varying positions across an IS3 backbone, and measured fluorescence from an adjacent reporter gene. We found that promoters need to occur near the end of the IS, within 150 bp from the reporter gene to drive detectable expression (**Fig S1**). This motivates our choice to study de-novo promoters emerging from the ends of IS3s.

We amplified five 150 bp sequences from the right (R) and left (L) ends of three *E.coli* IS3s. We call these sequences 1L, 1R, 2R, 3L, and 3R (1:IS150, 2:IS2, 3:IS3) (**Fig S3a**) (Methods). We chose these particular IS3s because they are well-characterized in *E. coli* and because they did not have detectable promoter activity. We pooled these *parent sequences* and created from them a deep mutational scanning library of *daughter sequences* via error-prone polymerase chain reaction (**Fig 1a**). We cloned the resulting library into the pMR1 plasmid to measure promoter activity on both DNA strands (top: GFP, bottom: RFP)^28^ (**Fig 1b**) and transformed the library into *E. coli*. We then carried out Sort-Seq^29–34^ (**Fig 1c**), separating bacteria with a cell sorter (BD Biosciences, FACSAria III) into eight fluorescence bins according to whether each daughter sequence drives no, low, medium, or high fluorescence for GFP and RFP. We sequenced the plasmid inserts from each subpopulation, and from the number of reads in each bin, calculated fluorescence scores (arbitrary units, a.u.) for the top and bottom strands, where 1.0 a.u. corresponds to no expression and 4.0 to strong expression (Methods).

**Figure 1.**
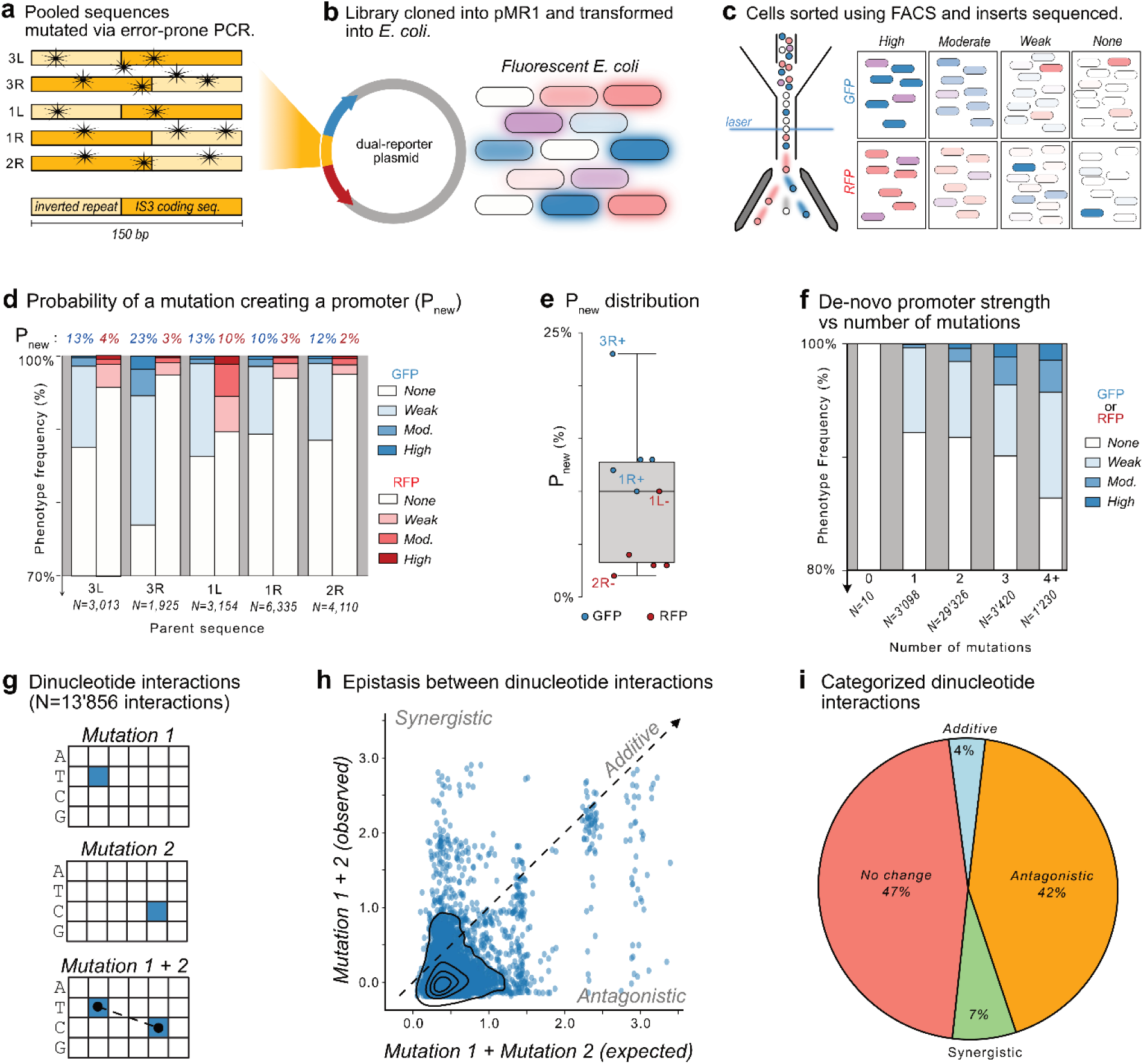
Promoters emerge with different probabilities for each parent sequence through epistatic interactions. **(a)** We pooled parental IS fragments for error-prone PCR, **(b)** cloned the mutagenized library into the dual reporter plasmid pMR1, and transformed the resulting plasmid library into *E. coli*. Inserts with promoter activity fluoresce green or red (shown as blue or red here), depending on the orientation of the newly created promoter, and with different intensities based on the promoter strength. **(c)** We sorted bacteria using fluorescence activated cell sorting (FACS) into four bins for each fluorescence color, corresponding to none, low, medium, and high fluorescence for both GFP and RFP (thus 8 bins total). We isolated inserts from cells in each bin and subjected them to Illumina sequencing. **(d)** The top of the figure shows for each parent sequence (x-axis), the probability of a mutation creating an active promoter de novo (Pnew) expressed as a percentage. The height of the vertical bars shows the percent of mutations creating promoters with expression strengths in each of four color-coded categories (color legend, blue: GFP, red: RFP). Note: the y-axis begins at 70%. **(e)** The probability of a mutation creating an active promoter de novo in the parent sequences (Pnew) for both the top strand (blue: GFP) and bottom strand (red: RFP). **(f)** Percent of de novo promoters in each strength category (white to blue, see color legend) based on the number of mutations. Note: the y-axis begins at 80%. **(g)** Schematic illustration of dinucleotide interactions based on double mutants where we also have mutational data on the individual mutants. The cartoon shows an example of two hypothetical individual mutations (upper and middle) and a double mutant (lower panel) at a given position (x-axis) and nucleotide (y-axis), where the blue square corresponds to the post-mutation nucleotide. There are 13,856 such interactions in our data. **(h)** For each dinucleotide interaction, we plot the fluorescence changes in arbitrary units (a.u.) for the sum of the individual mutations on the x-axis as “expected” values vs. the fluorescence change of their corresponding double mutant as an “observed” value. The dotted line indicates identity between expected and observed values and corresponds to additive interactions. Interactions above the dotted line are synergistic, where the observed expression is greater than expected from the sum, and those below are antagonistic, where the observed expression is less than expected. The black striations are kernel density estimates of the scatterplot (seaborn.kdeplot)^35^, and illustrate the central tendency of the data, which lies at −0.00 a.u. observed, and +0.41 a.u. expected. **(i)** We rounded the fluorescence changes of each dinucleotide interaction to the nearest 0.5 a.u. and classified the interactions as follows. If mut 1 = 0.0 a.u., mut 2 = 0.0 a.u., and mut 1+2 = 0.0 a.u., the interaction is classified as *no change.* If mut1 + mut2 = mut1+2, the interaction is classified as *additive.* If mut1 + mut2 > mut1+2, the interaction is classified as *synergistic.* If mut1 + mut2 < mut1+2, the interaction is classified as *antagonistic.* The percentages of the pie chart correspond to the ratio of each category for the 13,856 dinucleotide interactions.

We identified 18’537 unique daughter sequences, with a mean of ~3’707 daughters per parent (1L = 3’154, 1R = 6’335, 2R = 4’110, 3L = 3’013, 3R = 1’925), and a median of 2.0 point mutations per daughter (± 0.60) (**Fig S3**). Every parent and genetic orientation (top / bottom strand, 10/10) gave rise to daughter sequences with promoter activity, with 3’173 / 18’537 (~17%) daughters exhibiting promoter activity on either the top or bottom DNA strand (fluorescence score ≥ 1.5 a.u.). Of these, 2’408 / 18’537 (~13%) expressed GFP, 765 / 18’537 (~4%) expressed RFP (**Fig S3**), and 113 / 18,537 (~1%) expressed both GFP and RFP. For each parent sequence, we calculated the probability *P_new_* that mutation creates a new promoter (**Fig 1d,e**), which varies 11.5-fold among parent sequences (2R: *P_new_*=0.02, 3R: *P_new_*=0.23). Thus, relative to each other, some IS3 ends are biased towards evolving promoters, while others are biased against it (**Fig 1d,e**).

**Table 1.** Hotspot summary.

| Hotspot | Parent | Strand | Overlap | Mechanism | Figure |
| --- | --- | --- | --- | --- | --- |
| 1 | 1L | + | None | Unknown | 2d |
| 2 | 1L | + | None | Gain -35 box | S9l |
| 3 | 1L | - | -35 | Gain -10 box | 3b |
| 4 | 1L | - | -35 | Gain -10 box | 3c |
| 5 | 1R | + | -35 \| -10 | Gain -10 box | S9b |
| 6 | 1R | + | -35 | Gain -10 box \| Gain σ32 -10 box | S9c, S10b |
| 7 | 1R | - | -35 | Gain -10 box | S9h |
| 8 | 2R | + | None | Unknown | 2d |
| 9 | 3L | + | Promoter | Mutations flanking -35 box | 2d |
| 10 | 3L | + | Promoter | Mutations flanking -10 box | 2d |
| 11 | 3L | + | None | Gain σ28 -35 box | S10c |
| 12 | 3L | - | -10 or -35 | Gain -10 box | S9e |
| 13 | 3L | - | -35 | Gain -10 box | S9f |
| 14 | 3R | + | -35 | Gain -10 box \| -35 box | S9j, S8a |
| 15 | 3R | + | -35 \| -10 | -10 box | S8a |
| 16 | 3R | + | None | Gain σ28 -35 box \| Pausing site | S10d, S8 |
| 17 | 3R | + | None | Pausing site | S8 |
| 18 | 3R | - | -35 | Gain -10 box | S9n |

### The frequency of de novo promoters and their strength increases with further mutations

We next asked how the strength of de novo promoters relates to the number of mutations per daughter sequence. We categorized the daughter sequences by the number of mutations they harbor (1, 2, 3, and 4+), and determined promoter strengths for all daughter sequences in each category (none, weak, medium, and high) (**Fig 1f**). The frequency of daughter sequences encoding promoters increases with the number of mutations. Specifically, ~15% of all single mutants are promoters in either orientation (233 /1’549), along with ~16% of double mutants (2’343 / 14’663), ~19% of triple mutants (328 / 1’710), and ~25% of mutants with 4 or more mutations (156 / 615). For all daughters with single mutations, 85.0% have no expression on either strand (1’316 / 1’549), 14.5% convey weak expression (224 / 1’549), ~0.4% convey moderate expression (7 / 1’549), and ~0.1% convey high expression (2 / 1’549). (See **Fig S4** for genotype-phenotype maps). Conversely, for daughters with 4 or more mutations, ~75% have no expression on either strand (459 / 615), ~18% weak expression (111 / 615), ~5% moderate expression (28 / 615) and ~3% high expression (17 / 615). Thus, the frequency of promoter emergence and the frequency of strong de novo promoters increases with number of mutations.

### Antagonistic dinucleotide interactions in promoter emergence

Because ~15% of single mutants, but only ~16% of double mutants encode promoters (see Fig 1f), we hypothesized that individual mutations often interact non-additively (epistatically). To test this hypothesis, we identified all double mutants in our dataset, and retained only those whose constituent single mutations were also present in the dataset. The resulting 13’856 double mutants allowed us to identify dinucleotide interactions (**Fig 1g**). For each dinucleotide interaction, we plot the fluorescence changes of the double mutant (observed values) against the sum of the individual fluorescence changes (expected values) as a scatterplot (**Fig 1h**). An observed value identical to an expected value indicates additivity (no epistasis). An observed value greater than (less than) the expected value indicates synergistic (antagonistic) epistasis. The central tendency of the distributions lies below the 1:1 additive line (+0.41 a.u. expected, 0.0 a.u. observed), meaning that dinucleotide interactions are primarily antagonistic (**Fig 1h**). More specifically, most single mutations that increase promoter activity cancel out when combined (**Fig 1h**). Trinucleotide and tetranucleotide interactions are also mostly antagonistic (**Fig S5a,b**).

To categorize mutant interactions, we rounded the fluorescence changes to the nearest 0.5 a.u. for each dinucleotide interaction, and found that ~47% of double mutants exhibit no change in fluorescence (mut 1 = 0.0 a.u., mut 2 = 0.0 a.u., muts 1 + 2 = 0.0 a.u.), ~42% are antagonistic, ~7% are synergistic, and ~4% are additive (**Fig 1i**). Rounding to the nearest 0.25 a.u. increased the antagonistic category from ~42% to ~89% and lowered the no change category from ~47% to ~2% (**Fig S5c**). We also asked whether membership in each of these four epistatic categories is influenced by the distance between the mutated nucleotides, but did not observe a significant effect of distance on membership (ANOVA, p=0.584) (**Fig S5d**). In sum, most individual mutations that increase fluorescence interact antagonistically, i.e., combining them reduces their individual effects on fluorescence. This epistasis is independent of the distance between the individual mutations.

### Mutual information reveals emergence hotspots

To find out where promoters arise within the mutagenized IS3 parent sequences, we calculated, for each position *i* in each parent sequence, the mutual information *I_i_* between the identity of the nucleotide in each daughter sequence at this position (`A`,`T`,`C` or `G`), and the level of gene expression (fluorescence) associated with the nucleotide (1.0 – 4.0 arbitrary units, a.u.)^30–32^ **(Fig 2a)**. The essence of this calculation is to divide the joint probability of a position having a particular base *b and* fluorescence score *f*, *p_i_*(*b*, *f*), by the product of the individual probabilities *p_i_*(*b*) × *p*(*f*). This calculation can be visualized with a Venn-diagram, where the individual probabilities *p_i_*(*b*) and *p_i_*(*f*) are represented as circles, and the joint probability *p_i_*(*b*, *f*) corresponds to the area where the circles overlap **(Fig 2b)**. The higher the mutual information at position *i*, the more that position contributes to fluorescence changes, revealing where mutations create sites for transcription factor and σ factor binding (**Fig 2c**) (Methods).

**Figure 2.**
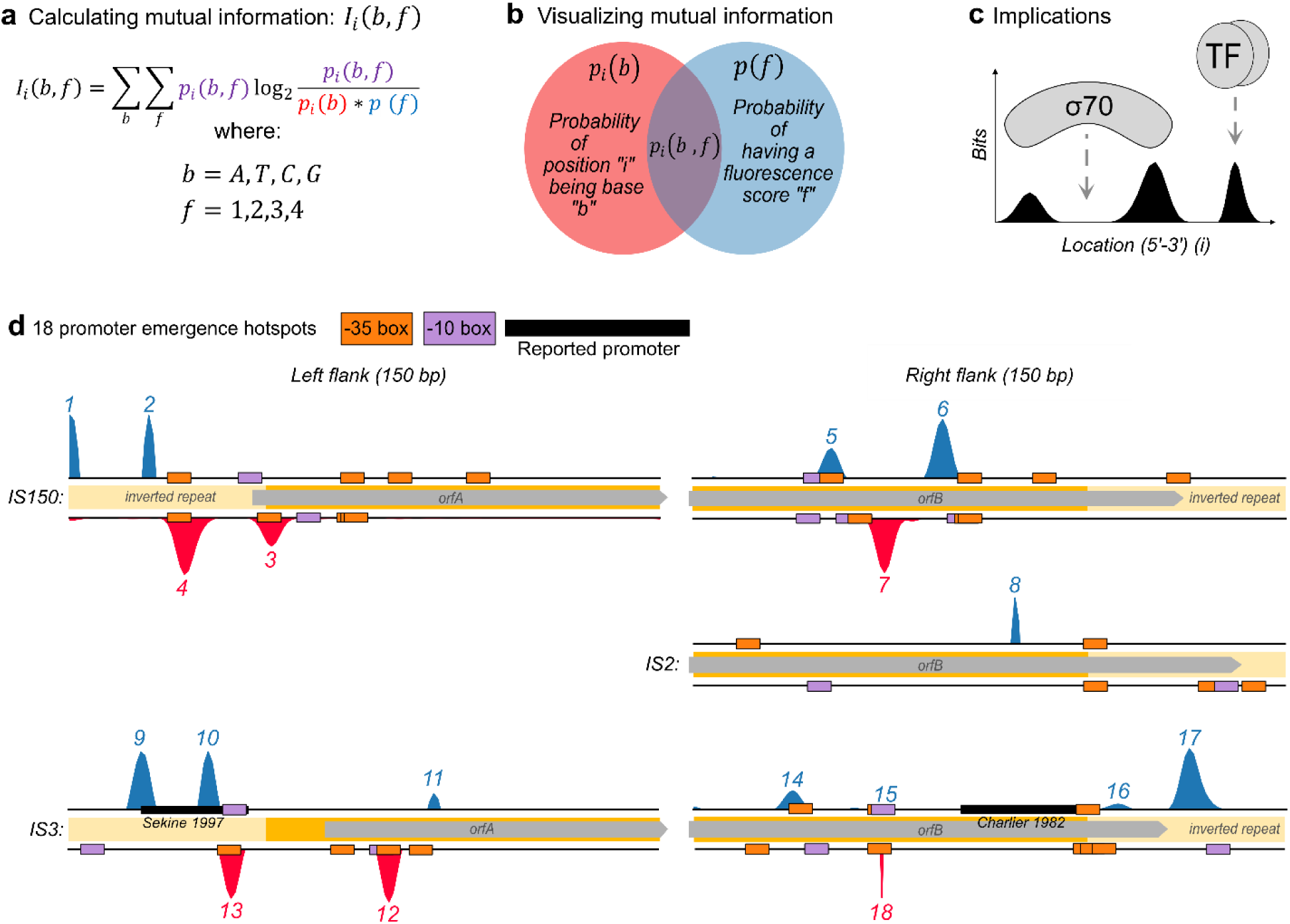
Mutual information reveals hotspots for de novo promoter emergence. **(a)** We calculated the mutual information *Ii(b,f)* for every position *(i)* in each parent DNA sequence, and for every possible base *(b = A, T, C, G)* and fluorescence value *(f=1,2,3,4)* with the equation shown here (see methods). **(b)** The components of the equation can be illustrated with a Venn-diagram, where the red circle corresponds to the probability *p_i_*(*b*) of a position i encoding base b, and the blue circle to the probability *p*(*f*) of being associated with a fluorescence score *(f).* The intersection of the two circles in magenta corresponds to the joint probability *p_i_*(*b*, *f*) of position *(i)* encoding base (*b*) and having a score of *(f)*. **(c)** Peaks of mutual information have been used in previous studies to map protein binding sites on DNA ^30,31^. **(d)** Mutual information for the five parent sequences in both top and bottom orientations. Mutual information reveals 18 regions (numbered histograms) in the parent sequences where daughter sequences are preferentially mutated to create fluorescence activity. We refer to these as promoter emergence hotspots. Orange and magenta boxes correspond to PWM-predicted −35 and −10 boxes present in the parent sequences. Black rectangles correspond to previously characterized promoters^3,16^. Light yellow areas correspond to an ISs inverted repeats, and gray arrows to open reading frames (orf) A and B. Note: to illustrate the location of the hotspots within the parent sequences, the y-axis scale differs among parent sequence. See **Figure S6** for a figure with identical y-axis scales.

We identified 18 regions of high mutual information which we call emergence *hotspots*^27^. Each parent sequence harbors 0-4 hotspots (see **Table 1**, **Fig 2d**, and **Fig S6**). Based on previous works studying promoter emergence and evolution^8,27^, we hypothesized that these hotspots correspond to 1) preexisting −10 and −35 boxes, 2) regions where new −10 and −35 boxes form, and 3) repressing sequences.

### Promoter activity occurs in regions with preexisting −10 and −35 boxes

To test whether the hotspots correspond to preexisting −10 and −35 boxes, we overlayed the mutual information with position-weight matrix (PWM) predictions for −10 and −35 boxes. PWMs are computational models based on protein-DNA binding experiments. When given a query sequence, they return a score predicting how strongly a focal protein binds to the query sequence^36^ (see methods). PWM analysis shows that the majority of hotspots (10/18) overlap with existing −10 or −35 boxes (see **Table 1** and **Fig 2d**). Additionally, hotspots #9 and #10 overlap with a previously characterized promoter in 3L(+) (**Fig 2d**).

This overlap suggests that some mutations which create promoters coincide with increasing −10 and −35 box PWM scores (see **Fig S4**). To validate this association, we calculated the changes in PWM scores of −10 and −35 boxes before and after each single point mutation (**Fig S7**). We found that −10 box PWM scores are ~1.5 times (14.2% vs 9.3%) more likely to increase when a single mutation creates a weak promoter than when it creates no promoter (chi-squared test, 4 d.f., p = 0.014). In contrast −35 box scores are no more likely to increase when a mutation creates a new promoter than when it does not (weak promoters: 12.8% vs promoter-neutral: 14.1%). Together, these findings suggest that promoters emerge when mutations create or modify −10 boxes, but not necessarily −35 boxes.

### Promoter activity emerges when mutations create new −10 and −35 boxes

We next hypothesized that the hotspots correspond to regions where new −10 and −35 boxes form. To this end, we computationally searched for regions in each parent sequence where mutations in a daughter gain a −10 or a −35 box, and tested whether the gain significantly associates with increased fluorescence (Methods). We found that ~50% (9/18) of the hotspots can be explained by mutations creating new −10 boxes, and one additional hotspot can be explained by gaining a −35 box (See **Fig 3a-d** and **Fig S9**. See also **Fig S10**, where we repeated this analysis using PWMs for the other sigma factors.

**Figure 3.**
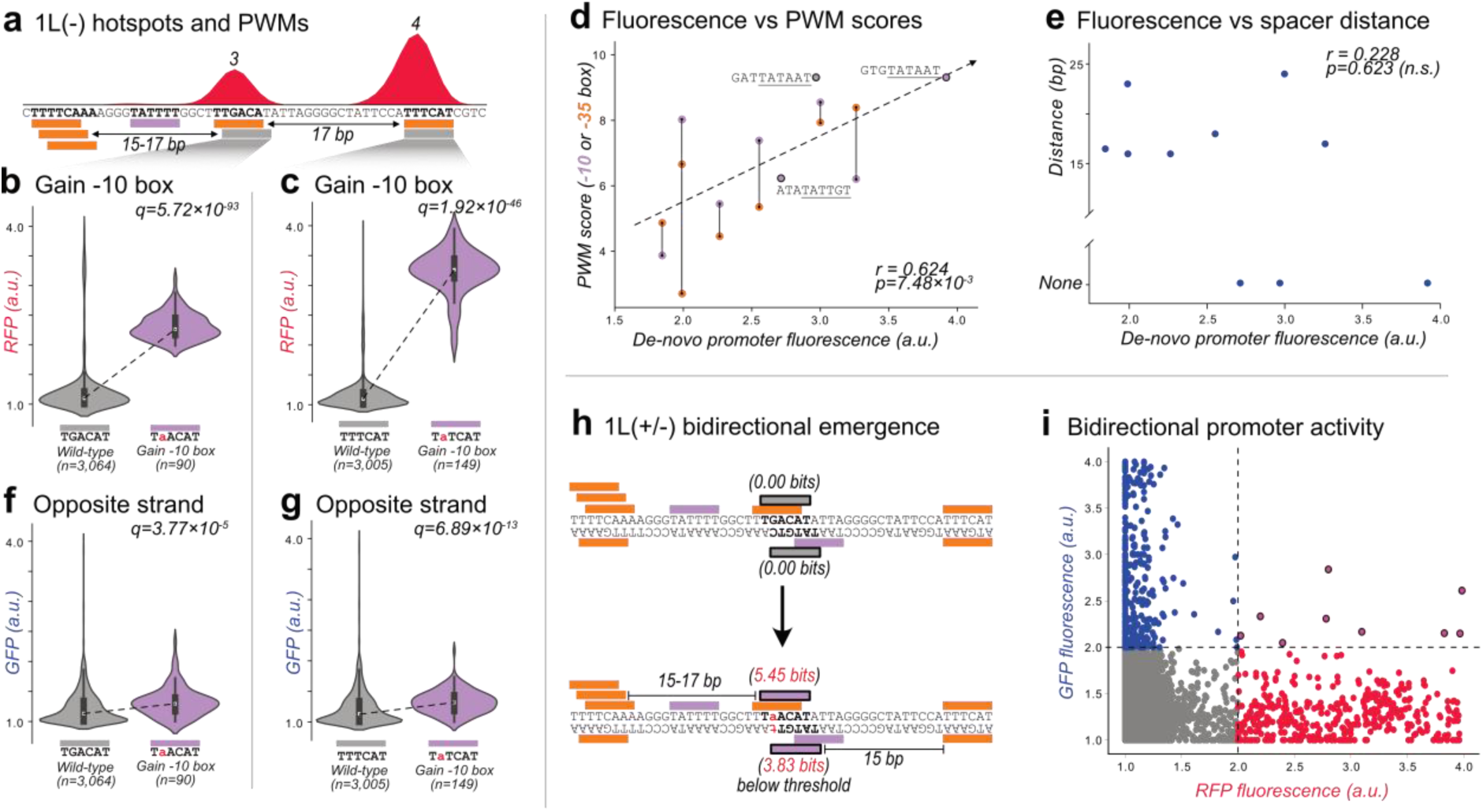
Unidirectional and bidirectional promoter emergence. **(a)** Parent 1L(−). Top: Mutual information between fluorescence scores and nucleotide identity at every position of the parent sequence. The numbers above each peak correspond to their “hotspot” identifiers (see Table 1). Bottom: Predicted −10 boxes (magenta rectangles) and −35 boxes (orange rectangles). Gray rectangles correspond to regions of interest. Bold sequences overlap with either a −10 or −35 box. Arrows indicate the distance between the regions of interest to the respective upstream −35 boxes. **(b)** RFP fluorescence scores for daughters without or with a −10 box in the region of interest shown in panel (a). The most frequent genotype is written below for each group. We test the null hypothesis that gaining a −10 box in this region of interest does not increase fluorescence (two-tailed Mann-Whitney U [MWU] test), and correct the p-values using a Benjamini-Hochberg correction as corresponding q-values (see methods, two-tailed MWU test, q=5.72×10-93). The area of each violin plot corresponds to the kernel density estimate of each distribution. Within each violin plot is a boxplot where the white circle corresponds to the median, the box to the interquartile range (IQR), and the whiskers to 1.5×IQR. **(c)** Analogous to (b) but for the other region of interest shown in (a) (two-tailed MWU test, q=1.92×10-46). **(d)** Position-weight matrix (PWM) scores of the −10 and −35 boxes for ten de-novo promoters (y-axis) plotted against their respective median fluorescence scores (x-axis, arbitrary units [a.u.]). We test the null hypothesis that there is no correlation between the PWM scores and the fluorescence scores using (Pearson’s r=0.624, p=7.48×10-3). **(e)** Analogous to (d) but for the distance between the −10 and −35 boxes (r=0.228, p=0.623, n.s. = non-significant). **(f)** Analogous to (b) but for the GFP fluorescence scores on the opposite strand (two-tailed MWU test, q=3.77×10-5). **(g)** Analogous to (c) but for the GFP fluorescence scores on the opposite strand (two-tailed MWU test, q=6.89×10-13). **(h)** PWM scores of the region of interest shown in (b) and (f) before (top) and after (bottom) a point mutation creates a bidirectional promoter. Orange rectangles correspond to −35 boxes, magenta rectangles to −10 boxes, and gray boxes with bold letters to the region of interest. **(i)** Red (x-axis) vs green (y-axis) fluorescence scores for all parents and daughter sequences,. Dotted lines separate fluorescence levels below and above 2.0 arbitrary units (a.u.). We additionally colored the daughter sequences depending on their fluorescence. Red: RFP ≥ 2.0. Blue: GFP ≥ 2.0. Magenta with black outlines: RFP and GFP ≥ 2.0.

We highlight two examples of how gaining a new −10 box can create promoter activity (hotspots #3 and #4) in 1L(−) (**Fig 3a**). First, at hotspot #3, 3’064 mutant daughter sequences do not encode a −10 box (consensus `TGACAT`) while 90 daughter sequences do (consensus `TaACAT`). The newly created −10 box is located 15-17 bp downstream of three overlapping −35 boxes. Gaining this −10 box is significantly associated with a doubling of gene expression (~106% fluorescence increase from 1.10 a.u. to 2.27 a.u., MWU test, q=5.72×10^−93^) (**Fig 3b**). Second, at hotspot #4, 3’005 daughter sequences do not encode a −10 box (consensus: `TTTCAT`), while 149 daughter sequences do (consensus `TaTCAT`). This −10 box is located 17 bp downstream of a −35 box. Gaining the −10 box almost triples gene expression (~196% increase in fluorescence from 1.10 a.u. to 3.26 a.u., MWU test, q=1.92×10^−46^) (**Fig 3c**).

We then asked how the strength of these de-novo promoters associates with their PWM scores and the spacing between the −10 and −35 boxes. From the ten hotspots associated with gaining new −10 boxes or a 35 box (**Fig 3a,b** and **Fig S9**), we also searched for a pre-existing box based on PWM predictions and their proximity. For 7 of 10 hotspots, an existing −35 or −10 box occurs within 25 bp from the new −10 box, while 3 of 10 hotspots did not have a −35 box within 25 bp of the newly created box. In addition, in the latter 3 hotspots, the new −10 boxes do not have an upstream `TGn` motif (**Fig 3d**), which can compensate for promoters without a −35 box^37^. The PWM scores of the −10 and −35 boxes for these promoters moderately correlate with their corresponding median fluorescence values (Pearson correlation, r=0.624, p=7.48×10^−3^) (**Fig 3d**). We did not find any association between the distances between boxes and the respective fluorescence values (Pearson correlation, r=0.228, p=0.628) (**Fig 3e**). In sum, these analyses suggest that PWM scores are a better predictor of de-novo promoter strength than spacer lengths.

### Gaining −10 boxes can create bidirectional promoters with unequal strengths

Hotspot #3 increases RFP expression by ~106% when gaining a −10 box on the corresponding (bottom) strand (see **Fig 3b**). Surprisingly, this gain also increases GFP expression on the opposite strand by ~14% (1.13→1.29 a.u., MWU test q=3.77×10^−5^) (**Fig 3f**). Similarly, gaining a −10 box at hotspot #4 increases RFP expression by ~196%, and GFP expression from the opposite strand by ~16% (1.13→1.32 a.u., MWU test, q=6.89×10^−13^) (**Fig 3g**). We asked whether other IS3 parent sequences also gain promoter activity on both strands when daughter sequences gain −10 boxes. This is the case for 3 of 9 parents in which gained −10 boxes lead to increased expression. In these sequences expression also increases on the opposite strand, albeit more weakly (**Fig S9**). Specifically, gaining a −10 box increases fluorescence on the same strand as the −10 box by 106%, 196%, 144%, and it increases fluorescence on the opposite by 14%, 16%, and 18%, respectively.

Bidirectional promoters can have a *symmetrical −10 box*, i.e., two overlapping −10 boxes on opposite strands (consensus: `TATWATA`)^38^. We asked if symmetrical −10 boxes could help to explain our findings about bidirectional promoters. Indeed, for 3 of the 9 new −10 boxes associated with de novo bidirectional activity, mutations indeed created motifs that either resemble −10 boxes (0.00 bits < PWM score < 3.98 bits), or that are −10 boxes (PWM score ≥ 3.98 bits) on the opposing strand. For example, mutations in parent 1L (see **Fig 3a,b,f**) create a −10 box on one strand (underlined, 5′-<u>TGACAT</u>A-3′ → 5′-<u>TaACAT</u>A-3′), which simultaneously creates a −10-like-box on the opposite strand (underlined, 3′-<u>TATGTC</u>A-5′ → 3′-<u>TATGTt</u>A-5′) (**Fig 3h**). This symmetrical −10 box lies 15-17 bp from −35 boxes on the appropriate strands. Symmetrical −10 boxes, however, are still insufficient to explain our findings. The reason is that 5 of 6 hotspots that do not create bidirectional promoters when gaining a −10 box also create symmetrical −10 boxes (see **Fig S12**). To summarize, symmetrical −10 boxes may contribute to bidirectionality, but are not sufficient to explain the origin of bidirectional promoters.

We then asked more generally how often mutations create bidirectional promoters (**Fig 3i**). A plot of green versus red fluorescence driven from the same daughter sequence shows an L-shaped distribution, with fluorescence primarily being either red or green. Emergent bidirectional promoters with expression ≥ 2.0 a.u. on both strands are rare (n=9, 3L: 2 sequences, 1L: 3, 2R: 3, 3R: 1, 1R: 0).

Thus far, we have shown that promoters emerge from either preexisting −10 and −35 boxes or by gaining new boxes upstream or downstream of preexisting boxes. When mutations create a −10 box, this can frequently create bidirectional promoter activity. Promoters also emerge when mutations create motifs for other sigma factors (see Fig S10). In addition, we found that hotspots #16 and #17 likely correspond to mutations that destroy a polymerase pausing site (see Fig S8 and Supplemental Results for extended analysis). See Table 1 for a summary of the hotspots.

### IS3s are enriched on their ends with DNA bearing promoter signatures

Previous reports have identified four IS3s with outward-directed promoters^13^, but it is unclear how frequent outward-directed promoters are among the more than 700 currently characterized IS3 sequences^2^. To find out, we estimated promoter locations in IS3 sequences using two different approaches.

First, we used PWMs to identify −10 and −35 boxes in 706 IS3s (**Fig 4a**). If we found a −10 and −35 box spaced 15-19 bp apart, we classified them as a *promoter signature* (Methods), which hints at but does not prove promoter activity. We found 2’026 such signatures, 1’263 of which (~62%) occurred on the bottom strand of DNA. To visualize the distribution of signatures, we partitioned the total length of each IS3 into 10 equidistant bins (median length 127±13 bp), and counted the signatures in each bin across all 706 IS3s (**Fig 4b**). On both strands, signatures are significantly non-uniformly distributed across the IS3s. The reason is that the distal-most 10 percent of IS3s contain more signatures on average (Kolmogorov-Smirnov (KS) test, top p=3.80×10^−5^, bottom p=1.82×10^−5^, Methods), with ~36% of all promoter signatures occurring on the top right, top left, bottom right, or bottom left ends (732 / 2’032).

**Figure 4.**
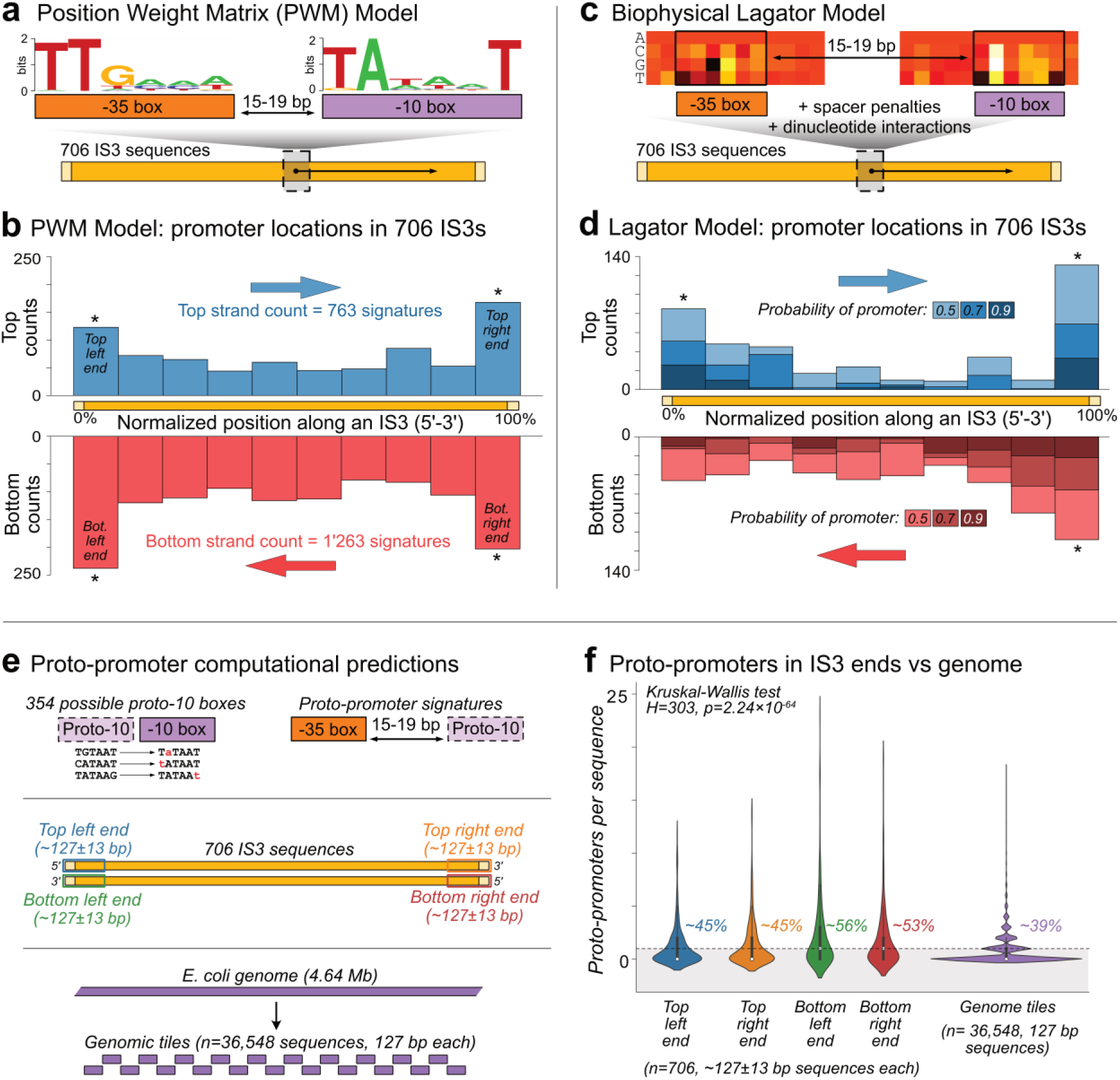
IS3s preferentially harbor promoter signatures close to their ends. **(a)** Sequence logos derived from position weight matrices (PWMs) depicting the likelihood of a base being bound at each position by a protein, in this case, the σ70 factor. The taller the letter at each position, the more likely it is that the corresponding nucleotide is present at the position in the binding site. Left: PWM logo for the −35 box. Right: PWM logo for the −10 box. To computationally identify promoter signatures in IS3s, we searched for −35 and −10 boxes using PWMs spaced 15-19 base pairs (bp) apart in both the top and the bottom strands of 706 IS3s. **(b)** We plotted the number of identified promoter signatures as histograms with a fixed bin width of 10% of IS3 length (10 bins in total). The top and bottom histograms correspond to promoter signatures on the top and bottom strand of IS3s, respectively. **(c)** The biophysical Lagator model uses energy matrices to identify −10 and −35 boxes (top left and right). The binding energies of σ70 to DNA sequences are represented as a heatmap, in which darker colors correspond to stronger binding (top left and right). The model additionally penalizes spacer distances which deviate from the canonical 17 bp, and accounts for dinucleotide interactions between and within boxes. **(d)** We divided each of the 706 IS3 sequences into 10 bins with a fixed bin width (10% of IS length), and predicted the number of promoters in each bin and sequence. Because the Lagator model only returns the probability of a query sequence being a promoter, we plot the number of promoters in each bin based on whether the predicted probability is greater than or equal to 0.5, 0.7, and 0.9, for both the top strand (blue histogram, top) and the bottom strand (red histogram, bottom). **(e)** Top left: proto −10 boxes are 6-mer DNA sequences that a −10 box PWM does not predict to be a −10 box, but that are one point mutation away from becoming one. Top right: proto-promoters are sequences that contain a proto −10 box with a −35 box 15-19 bp upstream. Middle: we look for proto-promoters in the first and final 10% of 706 IS3s on both the top and bottom strands. We refer to these areas as the top left (blue), top right (orange), bottom left (green), and bottom right (red) ends. The median length of an IS3 is 1,274 bp, with a standard deviation of 128 bp. Because the ends comprise 10% of each sequence, each end contains a sequence of 127±13 bp. Bottom: we computationally split the *E. coli* genome into 36,548 non-overlapping sequences that are each 127 bp long to match the length of the ends. We call each 127 bp sequence a genomic tile, and count the proto-promoters in them as well. **(f)** Number of proto-promoters per sequence in IS3 ends and genomic tiles. We test the null hypothesis that the central tendency of each distribution is the same, using a Kruskal-Wallis test (H=303, p=2.24 ×10-64).

Second, we identified putative promoters with a biophysical model we call the *Lagator model*^8^ (**Fig 4c**). Briefly, the Lagator Model uses binding energy matrices for −10 and −35 boxes that account for sequence composition outside of the core −10 and −35 box hexamers. The Lagator Model also incorporates dinucleotide interactions and penalizes sequences if their boxes are not canonically spaced 15-19 bp apart. Using the Lagator Model, we scanned 706 IS3s for promoter signatures, predicting the probability that each sequence and position encodes a promoter (see methods) (**Fig 4d**). While the Lagator Model predicts fewer promoters overall than PWMs, it still finds an enrichment of promoter signatures on the ends of IS3s (K.S. test at 50% probability, top p=4.40×10^−20^, bottom p=6.53×10^−21^, Methods).

We also validated our model predictions experimentally. To this end, we randomly selected five IS3s with and five IS3s without PWM-predicted outward-directed promoter signatures within 120 bp from their ends. We synthesized these terminal 120 bp, cloned them into our reporter plasmid, transformed it into *E.* coli (Methods), and measured fluorescence with a flow cytometer (BD Biosciences, FACSAria III). All tested IS3s (5/5) predicted to have outward-directed promoters drove higher fluorescence than the controls (**Fig S11**). Two of five IS3s not predicted to have outward-directed promoters did so as well (**Fig S11**). Extrapolating from this low false positive rate, we estimate that ~20% of putative promoter signatures (406 / 2’032) correspond to outward-directed promoters. These signatures occur on ~26% (181) of our 706 IS3s. In other words, we predict that at least a quarter of IS3s encode outward-directed promoters.

### The ends of IS3 sequences have more proto-promoter signatures on average than the genome

Approximately 50% of new promoters emerge from the ends of 10 IS3 sequences when point mutations create −10 boxes downstream of existing −35 boxes (Fig 3 and Fig S9). We asked whether we could predict such regions where promoters could arise through single mutations computationally. Specifically, we generated a set of all 6-mer DNA sequences that 1) were not classified as −10 boxes but 2) could become a −10 box with a single point mutation. We refer to these 354 sequences as *proto −10 boxes*. Next, we identified instances where a −35 box was located 15–19 bp upstream of a proto −10 box, defining these sequences as *proto-promoters* (**Fig 4e**).

Once having identified all proto-promoters, we asked if the 10% ends of IS3s contain more proto-promoters than the *E. coli* genome. To find out, we counted the number of proto-promoters on the top right, top left, bottom right, and bottom left 10% ends of IS3s (~127±13 bp each). In addition, we split the *E. coli* genome into 127 bp-long, non-overlapping sequences offset by one base pair (n=36,548), which we call *genomic tiles,* and counted the number of proto-promoters in them as well. We then plotted the distribution of proto-promoters per sequence for each sequence category (**Fig 4f**), and found a significant difference between the groups (Kruskal-Wallis test, H=303, p=2.24×10^−64^). Specifically, ~45% of IS3s have at least one proto-promoter on their top left end, ~45% on their top right end, ~56% on their bottom left end, and ~53% on their bottom right end. Among the genomic tiles, ~39% have at least one proto-promoter. Thus, the ends of IS3s harbor 1.15-1.44× more proto-promoters than the *E. coli* genome.

## Discussion

We created a mutagenesis library from the ends of five mobile IS3 sequences to explore how new promoters emerge in mobile DNA. We find that such non-regulatory sequences can evolve promoters through as little as one point mutation. Additional mutations can further increase the strengths of such de novo promoters. Most promoter-creating mutations occur in 18 hotspots. Approximately 50% of all promoters emerge from these hotspots when mutations alter or create new −10 boxes. IS3s harbor 1.15-1.44 times as many proto-promoter sequences compared to the *E. coli* genome. Motivated by this high latent regulatory potential, we also asked how likely it is that any IS3 encodes an outward directed promoter (without further mutation), and estimate that >26% (181 of 706) of IS3s do.

While most mobile DNA insertions are deleterious, an insertion can benefit the host if the mobile DNA drives fortuitous adjacent gene expression. Thus, mobile DNA that drives such expression may be preferentially preserved in evolution. For example, *E. coli* uses an IS3 as a mobile promoter to increase the expression of various genes during experimental evolution^16,39,40^. This has also been observed for other families of mobile DNA in pathogens, where mobile elements integrate at key genomic regions to ectopically express genes critical to conferring antibiotic resistances^41^. These findings are also consistent with the observation that eukaryotes use mobile DNA to evolve new cis-regulatory activity^13,14,42–49^. It is also worth noting that a subset of the IS3 sequences we studied are constrained from evolving new promoters. They have a low probability of evolving a promoter de novo and harbor no proto-promoters in their end regions. This lack of cis-regulatory potential in these IS3s may also influence evolution, since these IS3s would prevent the spontaneous emergence of new promoters or even abolish harmful gene expression.

All of the 10 IS3s and orientations we tested can evolve promoters from single mutations. This is important because single mutations provide the easiest route towards new promoters. Specifically, among 1,549 IS3 daughter sequences with single point mutations, ~7.8% acquired promoter activity. Additionally, the incidence of strong promoters increases with the number of mutations, such that sequences with four or more mutations are 24 times more likely to encode strong promoters than those with single mutations (1.44% vs 0.06%). We observed both synergistic and antagonistic epistatic interactions between these mutations. Thus, after mutation creates a new promoter, additional mutations can readily enhance that promoter’s activity without necessarily occurring in the core promoter sequence. Conversely, additional mutations can also readily remove a new promoter without necessarily occurring in the core. In other words, epistasis can allow selection to fine-tune the activity of new promoters.

A previous study in *E. coli* synthesized 40 randomly generated 103 bp parent sequences and tested their ability to evolve promoters de novo. ~10% (4/40) of the sequences already encoded promoter activity, and for ~60% (23/40), single point mutations sufficed to create promoters^24^. Another study used Sort-Seq and thermodynamic modeling to sample the genotypic space of 115 bp-long DNA sequences for promoters. It estimated that ~20% of such random sequences encode promoters, that ~80% can become promoters through single point mutations, and that ~1.5% of all single point mutations create promoters. For genomic DNA, these values were estimated to be lower^8^. By comparison, IS3s are 1.3-2.6 times more likely (26% vs 10-20%) to encode promoters than random sequences. In addition, 1.25-1.67 times as many IS3s (100% vs 60-80%) can evolve promoters from single mutations than random sequences. Moreover, ~15% of all single point mutations in IS3s create promoter activity on either DNA strand (~7.5% per strand), while in random sequences, only ~1.5% of mutations create promoter activity on a single strand^8^. Finally, IS3s contain 1.15-1.44 times as many proto-promoter sequences compared to the *E. coli* genome. Collectively, these findings suggest that IS3s may be a better substrate to create new promoters than both random and genomic DNA. This could potentially explain why IS3s persist in host genomes.

Mutual information has been used to map existing promoter architectures^30^. Here, we show that it can also map *future* cis-regulatory architectures. Specifically, mutual information helped us to identify 18 promoter emergence hotspots. We found likely explanations for 16 of these 18 hotspots. In 9 of the 18 hotspots, mutations create −10 boxes that increase expression, sometimes bidirectionally (3/9), and 1/18 create a −35 box (see Fig S9). The new promoters frequently emerge from hotspots when the new −10 box emerges downstream of a preexisting −35 box in regions we refer to as proto-promoters. 3/10 of the mapped de-novo promoters do not have a −35 box. This is consistent with estimates that ~20% of *E. coli* promoters do not encode −35 boxes^50^. Additionally, 12/18 hotspots overlap or are directly adjacent to preexisting −10 or −35 boxes, or previously described promoters^3,16^. Three hotspots also correlate with regions where mutations create motifs for other sigma factors (see Fig S10), and two hotspots in 3R(+) likely correspond to a polymerase pausing site (see Fig S8). The remaining 2 of 18 (~11%) hotspots have eluded characterization, but may overlap with IS3-specific transcription factor binding sites^13,51–53^, or with other unknown motifs awaiting discovery.

De novo promoters can be bidirectional, but almost exclusively drive stronger expression on one strand over the other. Only ~0.05% of daughter sequences (9/18,537) harbor bidirectional promoters with more than 2 a.u. of fluorescence from both strands. Given that ~19% of genomic *E. coli* promoters are bidirectional^38^, our finding suggests that such strong genomic bidirectional promoters are difficult to create de novo and that most bidirectional promoters exist due to positive selection for their bidirectionality.

Mobile DNA can be used by its host to evolve novel gene regulation^45–48^. In eukaryotes, many mobile DNA sequences have gene regulatory activity during development ^54^, and many young, species-specific regulatory elements (enhancers) originate from mobile DNA^42,45^. In prokaryotes, this has been observed in the evolution of antibiotic resistance^16,39–41,55^. The kind of mobile DNA we study here is well-suited for such co-option, because at least 26% of IS3s may already drive the expression of nearby genes. Those that do not can acquire this ability through as little as one point mutation. This latent potential for new gene regulation raises intriguing questions about its evolutionary origins.

## Acknowledgments

This work was supported by the European Research Council (Grant Agreement No. 739874), the Swiss National Science Foundation (grants 31003A_172887 and 310030_208174). TF is supported by a postdoctoral fellowship from the European Molecular Biology Organization (ALTF 963-2021) and a University of Zurich Postdoc Grant (FK-23-120). We thank all members of the Wagner group for discussions, Philipp Schätzle and Mario Wickert from the UZH cytometry facility for their training and support, and Baxter for enforcing a work-life balance.

## Supplemental Results

### A polymerase pausing site in 3R(+)

We hypothesized that promoter activity also emerges when mutations destroy repressor sequences. In particular within 3R(+), because we found a −35 box (`TTTAAT`, hotspot #14) and a −10 box spaced 15 bp apart (`TACAAT`, hotspot #15), which is indicative of an outward-directed promoter (**Fig S8a**). Additionally, a previous study described another outward-directed promoter in 3R(+) (`TTGGTG` and `CAATTT`)^16^, but this promoter does not overlap with any mutual information hotspot, nor does it strongly resemble a canonical promoter.

Downstream of these putative promoters are hotspots #16 and #17 (**Fig S8a**). We hypothesized that the DNA within these hotspots represses the promoters. To find out, we created a reporter construct without the last 50 bp of 3R(+), which excludes hotspots #16 and #17. Gene expression is indeed ~1.9× higher in this shorter construct (1,212 a.u. vs 638 a.u., two-tailed t-test, p=4.3×10^−188^), showing that the hotspot DNA represses transcription.

To find a candidate repressor protein, we tested the wild-type 3R(+) construct in a variety of genetic backgrounds that do not express the transcription factors FIS, IHF, and HNS, because these transcription factors can regulate ISs and their promoters^23,56–58^. We also tested 3R(+) in a genetic background without any Insertion Sequences^59^, because some studies found that IS3 OrfA proteins can repress promoters on ISs^3,52,60^. However, none of these experiments yielded a change in gene expression (**Fig S8b**).

Although repressor binding is thus not a likely cause for the transcriptional repression we observed, the repression could also be caused by a polymerase pausing site. Indeed a sequence overlapping with hotspot #16 and #17 resembles a consensus pausing site sequence^61^ (**Fig S8c**). In addition, mutations in key nucleotides of this sequence increase fluorescence (**Fig S8d,e**). Moreover, a pausing site can terminate transcription if it occurs downstream of a GC-rich Rho-utilization site (rut)^62,63^, which we also find in 3R(+) (**Fig S8a**). Such terminator sequences have also been described on the end regions of other insertion sequences^64,65^. Based on this information, we conclude that 3R(+) probably gains promoter activity when mutations destroy a polymerase pausing site.

## Materials and Methods

### Data availability

Python scripts, an anaconda environment with relevant packages and their versions, and supplemental tables can be found on Github: https://github.com/tfuqua95/promoter_emergence_mobile_DNA

Raw sequencing reads are accessible from the Sequence Read Archive (SRA) with the accession number (PRJNA1021969): https://www.ncbi.nlm.nih.gov/bioproject/PRJNA1021969

**Data S1 (separate file).** An Excel spreadsheet with a list of primers and DNA sequences to recreate all of the constructs tested in this manuscript.

**Data S2 (separate file).** A large csv file with each unique daughter sequence, its respective parent sequence, and the GFP and RFP fluorescence scores from the sort-seq experiment.

**Data S3 (separate file).** Contains a large csv file with the regions of interest and their respective associations with gaining −10 or −35 boxes and changing fluorescence. The table additionally includes the raw p-values and the corrected q-values.

**Data S4 (separate file).** Contains an Excel sheet with the data to rapidly reproduce the main figures in the text.

### Bacterial strains

We tested the reporter constructs in *E. coli* DH5α electrocompetent cells (Takara, Japan, product #9027). In the experiments of **Fig S8b**, we tested 3R(+) constructs in the following backgrounds. The *Δfis*, *Δihf*, and *Δhns* are from the KEIO collection^57^ (*Δfis*: *JW3229*, *ΔihfA: JW1702*, *Δhns*: *JW1225*). The *ΔIS* strain is derived from ref^59^ (Scarab Genomics, Clean Genome ® E-6265-05, USA).

### DNA sequences

We acquired al IS3 sequences from the ISfinder database^2^ (https://isfinder.biotoul.fr). See **Data S1** for a list of primers and DNA sequences.

### Molecular Cloning

To 1) linearize the pMR1 plasmid (cutting); 2) amplify DNA synthesized by Integrated DNA Technologies (IDT), USA); and 3) amplify the inserts from the *E. coli* genome, we used a high-fidelity Q5 polymerase (NEB, USA product #M0491). For each polymerase chain reaction (PCR), we added 1 uL of each primer at a concentration of 100 uMol, 5 uL of the provided Q5 reaction buffer, 1uL of template DNA, 1 uL 10 mM dNTPs (Thermo Scientific, USA, product #R0191), 1 uL of Q5 polymerase, and molecular grade water (AppliChem, Germany, product #A7398) to a volume of 50 uL per reaction. In a thermal cycler (C1000 Touch Thermal Cycler, Bio-Rad, USA) we performed each PCR for 30 cycles, annealing at 55°C for 30 seconds, and extending for 30 seconds at 72°C. See **Data S1** for a list of primers and DNA sequences. We separated the PCR products by size using gel electrophoresis, isolating the band of interest with a scalpel, and purifying the product with a Qiagen QIAquick Gel Purification Kit (Qiagen, Germany, product #28706). We carried out the gel purification according to the manufacturer’s instructions, apart from the final elution step, where we eluted all samples with 30 uL of H_2_0 instead of 50 uL of Elution Buffer to increase the DNA concentration. We estimated the concentrations of each purified product using a Nanodrop One spectrophotometer (Thermo Scientific, USA).

We cloned the inserts for measuring reporter activity in this study into the pMR1 dual reporter plasmid ^28^ between the EcoRI (`GAATTC`) and the BamHI (`GGATCC`) restriction sites. To clone the inserts, we used the NEBuilder kit (New England Biolabs [NEB], USA, product #E2621). Specifically, in a PCR tube, we added 100 ng of linearized pMR1 plasmid (plasmid linearized via PCR), 25 ng of the insert, 5 uL of the provided NEBuilder mastermix, and molecular grade water (AppliChem, Germany, product #A7398) to a volume of 10 uL. We incubated the reaction for 1 hour at 50°C in a thermal cycler (Bio-Rad C1000 Touch Thermal Cycler, Bio-Rad, USA).

We immediately transformed the cloned products into *E. coli* DH5α electrocompetent cells (Takara, Japan, product #9027), adding 2 uL of the product to 100 uL of electrocompetent cells. We then electroporated the cells with a Bio-Rad MicroPulser (Bio-Rad, USA) and 2mm electroporation cuvettes (Cell Projects, England, product #EP-202). We allowed the transformed bacteria to recover in 1 mL of the “Super Optimal Broth with Catabolite Repression Medium” (SOC) medium provided with the electrocompetent cells, and incubated the bacteria at 37°C, shaking at 230 RPM for 1.5 hours (Infors HT, Switzerland, Multitron). After the incubation, we plated 5 uL of the bacteria onto a standard petri dish using glass beads on LB-Agar medium supplemented with 100 ug/ml of chloramphenicol. With the remaining ~995 uL of the bacteria culture, we transferred the bacteria to a 50 mL tube, and added 9 uL of LB-chloramphenicol (100 ug/ml) for a total volume of ~10 uL. We incubated the culture overnight at 37°C shaking at 230 rpm. The following morning, we combined 1 mL of the culture with 667 uL of 60% (weight / volume) glycerol, and stored the library at −80°C until needed. To verify the sequence of a cloned insert, we randomly selected three colonies from the LB-agar plate and sequenced using Sanger sequencing (MicroSynth, Switzerland).

### Control sequences

We created three control plasmids to identify confounding factors contributing to IS-driven gene expression through fluorescence activated cell sorting (FACS, see **Fig S2c and S3b,c**). The first is a GFP-positive control, for which we cloned the bba_j23110 promoter oriented towards the GFP coding sequence of pMR1. The second is an RFP-positive control, which also harbors the bba_J23110 promoter, but we cloned it in the opposite direction to face the RFP coding sequence of pMR1. The third control is an empty pMR1 plasmid without an insert between the BamHI and EcoRI cut sites. We cloned these inserts and transformed the products as described in the “Molecular Cloning” section.

### Cytometry plots

We analyzed the flow cytometry data from .fcs files using the software FlowCal ^66^. We prepared all plots using the python libraries seaborn ^35^ and matplotlib ^67^. Data and software version numbers are available on the GitHub repository: https://github.com/tfuqua95/promoter_emergence_mobile_DNA

### Error-prone PCR

To create the mutagenesis library, we prepared a 100 uL GoTaq (Promega, USA, product #M3001) polymerase chain reaction (PCR). For this reaction, we added 1 uL of the forward and reverse primer at a concentration of 100 uMol, 20 uL of GoTaq reaction buffer, 1uL of template DNA, 1 uL 10 mM dNTPs (Thermo Scientific, USA, product #R0191), 1 uL of GoTaq polymerase, 1 uL of 15 mMol MnCl_2_, and molecular grade water (AppliChem, Germany, product #A7398) to a volume of 50 uL per reaction. For the template DNA, we combined an equimolar ratio of each parent sequence. The supplemented MnCl_2_ provides the mutations. See **Data S1** for a list of primers and DNA sequences.

In a thermal cycler (C1000 Touch Thermal Cycler, Bio-Rad, USA) we performed each PCR for 30 cycles, annealing at 55°C for 30 seconds, and extending for 30 seconds at 72°C. We separated the PCR products by size using gel electrophoresis, selecting the band of interest with a scalpel, and purifying the product with a Qiagen QIAquick Gel Purification Kit (Qiagen, Netherlands, product #28706) according to the manufacturer’s instructions. We only deviated from the protocol at the final elution step, where we eluted all samples with 30 uL of H_2_O instead of 50 uL of TE buffer. We verified the concentrations of each purified product using a Nanodrop One spectrophotometer (Thermo Scientific, USA). We then cleaned the product and transformed it into *E. coli*, as described in “Molecular Cloning”.

Because we pooled the template sequences at the beginning of the reaction, the library contained different amounts of mutant daughter sequences for each parent template sequence (**Figure S3d**). Because of this amplification bias, we excluded the parent sequence 2L from the analysis in this study. For future studies, we recommend carrying out individual error-prone PCR reactions per parent sequence, and then pooling the products after purification.

### Fluorescence activated cell sorting (FACS)

We inoculated 100 uL of the error-prone PCR library glycerol stock (see sections “Error-prone PCR” and “Molecular Cloning”) into a 1 mL LB-chloramphenicol solution (100 ug/ml chloramphenicol), and let the resulting culture grow overnight at 37°C, with shaking at 230 rpm (Infors HT, Switzerland, Multitron). The following morning, we washed the culture twice in Dulbeco’s Phosphate Buffered Saline (PBS) (Sigma, USA, D8537) before sorting cells with an Aria III fluorescence activated cell sorter (BD Biosciences, USA) into eight fluorescence bins (GFP and RFP: none, low, medium, and high). To detect and measure GFP fluorescence, we used a 488 nm laser, measuring fluorescein height (FITC-H) at 750 volts. For RFP, we used a 633 nm laser, measuring phycoerythrin height (PE-H) at 510 volts.

To draw the fluorescence gates, we defined fluorescence bin boundaries based on fluorescence measurements from the following three control plasmids. GFP-control: bba j23110 promoter oriented towards GFP. RFP-control: bba j23110 promoter oriented towards RFP. Negative control: empty pMR1 plasmid. See also **Figure S3b,c** and “Control Sequences”.

We define the lower boundary of bin #1 (none, i.e. no expression) for green fluorescence, as the minimum of (i) the lowest value of measured green fluorescence for the negative control (empty pMR1) and (ii) the lowest value of measured green fluorescence for the positive control, but for the opposing fluorophore (RFP). We define a lower boundary for the lowest fluorescence bin to prevent artefacts that may arise when a cell sorter sorts various debris into the lowest bin, including but not limited to salts, empty droplets, or bacterial waste. We define analogously the upper boundary of bin #1 as the maximum of (i) the highest value of measured green fluorescence for the negative control (empty pMR1) and (ii) the highest value of measured green fluorescence for the positive control, but for the opposing fluorophore (RFP). We define the lower and higher boundaries of bin #1 for red fluorescence analogously, but with switched roles for GFP and RFP.

We defined the lower boundary of bin #4 (high, i.e., highest expression) as the mean fluorescence of the respective (green or red) positive control. Because this was the bin with the highest fluorescence, we did not define an upper bound for bin #4.

To define bins #2 and #3, we divided the interval between the lower boundary of bin #4 and the upper boundary of bin #1 in half, and set the upper bound of bin #2 and the lower bound of bin #3 to this half-way point. See **Figure S2c and S3b,c** for the division of all bins.

We sorted the mutagenesis library over two consecutive days. After sorting at the end of the first day, we added 1mL of SOC medium (Sigma, USA, product #CMR002K) without antibiotics to the sorted cultures, and let the cells recover for two hours at 37°C, with shaking at 230 rpm (Infors HT, Switzerland, Multitron). Afterwards, we filled the cultures with LB-Chloramphenicol (100 ug/ml chloramphenicol) to 10 mL and let the cultures grow overnight, incubating and shaking them as just described.

To ensure that we had sorted each genotype into the appropriate fluorescence bin, we repeated the sorting on the following day using the same procedure. For example, if we had sorted cells that fluoresce at low levels into bin #2 on the first day, we sorted daughter cells from this culture on the second day only into bin #2, i.e., allowing only cells whose fluorescence falls into the boundaries of this bin to be considered for the next analysis step (DNA sequencing). This re-sorting step ensures that we only sequence genotypes that are sorted into the same fluorescence bin after both consecutive days, lowering the possibility of sorting errors. To further minimize these sorting errors and to estimate the variance in fluorescence levels, we also sorted cells into three technical triplicates (r1, r2, r3, see **Fig S3g,h**) on the second day. In the context of the example, this means that on the second day, we sorted the culture from bin #2 into bin #2 three times, i.e., in three replicate sorting experiments (r1-2, r2-2, r3-2). **Table S1** describes the number of cells and replicates sorted into each bin.

After the second round of sorting, we once again allowed cells to recover in SOC and grew the cultures overnight, as previously described for day 1. The following morning, we created a glycerol stock by adding 1 mL of the culture and 667 uL of 60% glycerol (weight by volume) to a cryotube and stored the cultures at −80°C. We prepared the remaining culture for DNA isolation and Illumina sequencing (see Illumina Sequencing).

To summarize, from a single mutagenesis library of bacterial cells, we sorted bacteria into 24 individual cultures, where 12 cultures correspond to green-sorted bins (GFP) and the other 12 to red-sorted bins (RFP). For both green and red fluorescence, we sorted cultures into three replicates (r1, r2, and r3), each of which we binned into four fluorescence levels (none, low, medium, and high, corresponding to bin#1, #2, #3, and #4 respectively).

### Illumina Sequencing

From each sorted culture (see “Fluorescence activated cell sorting” section), we isolated plasmids using a Qiagen QIAprep Spin Miniprep Kit (Qiagen, Germany, product #27104), following the manufacturer’s instructions apart from eluting the DNA in 30 uL of H_2_O instead of 50 uL of Elution Buffer. From the isolated plasmids, we PCR-amplified the plasmids’ inserts using Q5 polymerase (NEB, USA product #M0491) (see “Molecular Cloning” for protocol). We multiplexed the forward primer for each PCR with a unique barcode for each bin and replicate (r1-bin1-GFP, r2-bin1-GFP, r3-bin1-GFP, r1-bin2-GFP, …, r3-bin4-RFP.). In addition, we also isolated plasmids from the unsorted library and PCR-amplified their inserts with their own unique barcoded primers (24 + 1 = 25 total PCRs). See **Data S1** for a list of primers and barcodes.

We separated the resulting PCR products by size using gel electrophoresis, selecting the band of interest using a scalpel, and purifying the product with a Qiagen QIAquick Gel Purification Kit (Qiagen, Netherlands, product #28706) according to the manufacturer’s instructions. We only deviated from these instructions in the final elution step, where we eluted all samples with 30 uL instead of 50 uL of the provided elution buffer. We verified the concentrations of each purified product using a Nanodrop One spectrophotometer (Thermo Scientific, USA). We then pooled the barcoded samples and sent them for Illumina paired-end read sequencing (Eurofins GmbH, Germany).

### Processing sequencing results

We merged paired-end reads using Flash2^68^. Paired-end reads can be sequenced in either genetic orientation, which can result in ambiguous read orientations. To avoid such ambiguities, we took advantage of the fact that all our inserts were cloned between the palindromic 5’-EcoRI (`GAATTC`) and 3’-BamHI (`GGATCC`) restriction sites of pMR1. We searched for both sites in each paired-end read and discarded any paired-end reads that did not encode both sites. If the BamHI site was upstream of the EcoRI site, we used the reverse complement of the paired-end read for further analysis. The result was that all the paired-end reads are in the same orientation and contain both restriction sites. We then searched for the barcode upstream of the EcoRI site in each paired-end read, used it to identify the bin from which the read originated, and cropped the EcoRI and BamHI sites from each read. We counted the number of reads within each bin, and then created a table in which the first column contains a list of unique sequences. Further columns contain the number of reads associated with the unique sequences in different fluorescence bin (**Data S2**). We henceforth refer to each unique paired-end read as a “daughter sequence.”

We next removed any daughter sequence with a length different from 150 bp to focus on point mutations rather than insertions and deletions during the analysis. For each daughter sequence, we then calculated the Hamming Distance between the daughter sequence and each of the wild-type “parent sequences”, i.e., the number of nucleotide differences between these sequences. We assigned the daughter sequence to the parent sequence with the lowest Hamming Distance.

We determined fluorescence scores that indicate how strongly each daughter sequence drives the expression of RFP and GFP. To this end, we first calculated a fluorescent score (*F_rep_*) for each of our three technical replicates (r1, r2, and r3) with equation (1):

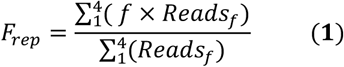

In this equation, *f* corresponds to the different fluorescence bins (none, low, medium, and high), which we integer-encoded as *f* = 1, 2, 3, 4, respectively. *Reads_f_* corresponds to the number of reads within each fluorescence bin *f*. As an example, **Table S2** shows the number of read counts of a specific sequence in each bin of replicate r1, which yields a final green fluorescence score of *F_r_*_1_=(1×49) + (2×4) + (3×3) + (4×0) / (49 + 4 + 3 + 0)=1.179 arbitrary units (a.u.) of fluorescence.

We calculated *F_rep_* for each technical replicate and each sequence, and averaged these replicate scores to compute a final fluorescence score. In addition to sequences and read counts, **Data S2** also provides these scores. We additionally calculated the standard deviation between the three replicates, and compared the fluorescence scores among replicates using a Pearson correlation coefficient (see **Fig S3g,h**).

We next filtered our data for quality control, removing daughter sequences from further data analysis that 1) are not also found in the unsorted library; 2) did not have at least one read in each of the replicates (r1, r2, r3); 3) are matched to a parent sequence with a Hamming distance larger than 10; 4) have a total number of fewer than 10 reads in all bins; 5) have a standard deviation between the three replicate fluorescence scores *F_rep_* greater than 0.3.

After this filtering step, 18,537 unique daughter sequences remained for further analysis, with a mean of 3,707 daughter sequences per parent sequence (3L = 3,013, 3R=1,925, 1L = 3,154, 1R = 6,335, 2R = 4,110). See also **Figure S3** for pertinent data.

### Position weight matrices (PWMs)

We obtained the PWMs for the −10 and −35 sites as a list of −10 and −35 sequences from Regulon DB ^69^. We converted the list of −10 and −35 sequences into a PWM using the `Biopython.motifs` package ^70^. To calculate a PWM, we needed to provide a background nucleotide composition. Because we aimed to use the PWMs for many different kinds of sequences, we set this background composition to equal 25% each for `A, T, C, and G.`

From a query sequence, a PWM returns a score in bits. The higher the score is, the higher the likelihood is that the query sequence binds the protein of interest. Because PWM scores can vary widely among different query sequences, it is not always clear when a PWM score is high enough that the query can be classified as a bona fide transcription factor binding motif. In our study, unless otherwise specified, we used the well-established “Patser threshold” for this purpose, which equals the information content of a motif ^36^. For PWMs used in this study, the −35 box has an information content of 3.39 bits, and for the −10 box 3.98 bits. We classified query sequences with a score greater than or equal to these thresholds as binding motifs.

When searching for promoter signatures in 706 IS3s, we first searched for −35 boxes using the −35 box PWM and the `motifs.search` function in Biopython ^70^. The function identifies both the location and score of all motifs above the specified threshold in the query sequence. If we found a −35 motif, we then searched for −10 boxes downstream of the −35-motif, using the −10 box PWM. If the sequence also encoded a sufficiently high-scoring −10 motif 14-17 downstream of the −35 motif, we classified the sequence as having a promoter signature.

To calculate how PWM scores both the −10 and −35 boxes change in response to single mutations, we first calculated the total PWM scores for both −35 and −10 boxes in the wild-type parent sequences. We then isolated a list of daughter sequences with single mutations that created weak promoter activity (**Fig. S7**), and a list of daughter sequences with single mutations that did not create promoter activity. For each subset of sequences, we calculated the PWM scores again. We then quantified the differences in the scores before and after the mutation, and created the contingency tables in **Figure S7**, classifying a mutation as either increasing, decreasing, or not changing the PWM score for both the −10 and −35 boxes. Because we were calculating the differences in scores, and not necessarily looking for the gain or loss of binding sites, we lowered the PWM threshold values for the −35 box (3.39 bits) and the −10 box (3.98 bits) to 0.00 bits each while searching for motifs.

### Identifying promoter signatures with the Lagator model

For a given query sequence, we first computationally extracted the first 36 bps of the sequence. We then computationally added 40 G’s upstream and downstream of this extracted sequence. We calculated the probability that the resulting 116 bp sequence has promoter activity (P_on_) using the “Extended Model” from ref ^8^, which we refer to as the Lagator Model. We then repeated this procedure, sliding a 36bp long window in +1 bp increments across the query to extract 36bp subsequences, and recalculated P_on_ for each subsequence. This computation results in an array of promoter probabilities, which when plotted along the length of the original sequence, reveals peaks, i.e., regions where the Lagator model predicts a promoter’s location. We identified these peaks using `find_peaks` from `scipy.signal`, which returns the position of peaks above a height of either 0.5, 0.7, and 0.9, numbers that refer to the respective probabilities that a promoter is present in Figure 4d. We performed this sliding window analysis for each of the 706 IS3s. To plot the locations of the promoter signatures as a histogram, we normalized the positions of the identified peaks to the length of the respective sequences from which the peaks are derived (see Fig 4d).

### Association between the gain and loss of −10/−35 boxes and fluorescence changes

For the analyses of **Figure 3** and **Figure S9**, we computationally searched for regions in each parent sequence that gained or lost −10 and −35 boxes through mutations that are also associated with significant fluorescence increases.

To search for these regions, we moved a sliding window of length 6 bp through the parent sequence (−10 and −35 boxes have a length of 6 base pairs). Within this window, we searched for either a −10 or −35 box motif in all of the parents’ mutant daughter sequences, as described in “Position Weight Matrices”. If the sequences in the sliding window contained a −35 or a −10 motif above the Patser Threshold ^36^ (−35 box = 3.39 bits, −10 box = 3.98 bits), we added the fluorescence scores to a list of motif “positives”, and otherwise to a list of motif “negatives”. If each list contained more than 10 fluorescence scores, we tested the null hypothesis that the two lists had the same fluorescence scores, using a one-sided Mann-Whitney U test with the `mannwhitneyu` function from `scipy.stats`.

We repeated these procedures for all positions of the sliding window within the parent sequence, from the beginning (position 1) to the end (position 150-6=144). We performed this analysis on all five parent sequences, both on the top and bottom strands, for −10 and −35 box motifs, and for both green and red fluorescence scores. Because we thus performed multiple hypothesis tests, we corrected all of our p-values into q-values using a Benjamini-Hochberg correction (false discovery rate = 0.05) ^71^. We classified a region as significantly associated with a gain in promoter activity when the test rejected the null hypothesis at q<0.05.

To focus our analysis on mutations with large effects sizes, we only report fluorescence gains greater than 10% that also partially overlap with the emergence hotspots in the manuscript. **Data S3** provides all of the identified significant changes, along with a list of the p-values and corrected q-values.

For hotspot #12 (**Fig S9e**), we hypothesized that mutations that increase the PWM score of an existing −10 box are associated with increased promoter activity. We tested this hypothesis by tabulating the fluorescence scores of (i) daughters with a −10 box PWM score less than or equal to the wild-type score, and (ii) daughters with a −10 box PWM score greater than the wild-type score. We then tested the null hypothesis that the two categories of daughters had the same fluorescence scores using a one-sided Mann-Whitney U test with the `mannwhitneyu` function from `scipy.stats`. Because this analysis was done post-hoc, we do not report a q-value.

For hotspot #7 (**Fig S9h**), we hypothesized that mutations create a weak −10 box that was below our threshold limit of 3.98 bits. We tested this hypothesis by tabulating the fluorescence scores of (i) daughters with a PWM score of 0.00 bits at the given position, and (ii) daughters with a PWM score greater than 0.00 bits (but less than 3.98 bits). We then tested the null hypothesis that the two groups of daughters had the same fluorescence scores using a Mann-Whitney U test. Because this analysis was done post-hoc, we do not report a q-value.

For hotspots #2, #14, and #18 (**Fig S9j,l**), we hypothesized that mutations create a −10 box and a −35 box, respectively, but our analysis could not detect this because the boxes were created in fewer than 10 daughter sequences. We grouped daughters with and without the box of interest, but did not have the necessary sample sizes to carry out a Mann-Whitney U test. For this reason, we do not provide a p or q-value.

### Testing additional sigma factor binding motifs

We acquired position weight matrices (PWMs) for additional sigma (σ) factors. Specifically, we acquired the σ_32_ −35 and −10 box PWMs from ref^72^, the σ_H_ −35 and −10 box PWMs from ref^73^, the σ_28_ −35 and −10 boxes from ref^74^, and the σ_54_ from ref^75^. Logos were drawn using Logomaker^76^. We then repeated the analysis described in the subsection: *association between the gain and loss of −10/−35 boxes and fluorescence changes*.

### Mutual information

Mutual information is a measure of dependence between two variables. We calculated the mutual information *I_i_* between the nucleotide identity *b* at position *i* of daughter sequences of a given parent (*1≤i≤150*), and the fluorescence score *f* for daughter sequences of a given parent. To calculate the mutual information for each parent sequence, we used equation (2) as previously described in ^31^:

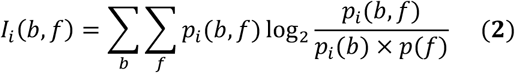

In this equation, the variable *b* represents all possible nucleotides (*b* = *A*, *T*, *C*, *G*). The variable *f* represents fluorescence scores rounded to the nearest integer (*f* = 1,2,3,4) (see “Processing sequencing results” for calculation of these scores); *p_i_*(*b*) corresponds to the probability (relative frequency) of each sequence variant encoding an `A, T, C,` or `G` at position (*i)*; *p*(*f*) corresponds to the probability (relative frequency) of fluorescence scores being equal to 1, 2, 3, or 4; and *p_i_*(*b*, *f*) is the corresponding joint probability, i.e., the probability of position *i* encoding an `A, T, C,` or `G`, <u>and</u> having a fluorescence score of 1, 2, 3, or 4 a.u.

The concept of mutual information is best illustrated with two simple examples. For the first, we calculate the mutual information for two consecutive and fair coin flips. Here, *b* equals the possible states of the first coin flip (heads or tails), and *f* equals the possible states of the second coin flip (heads or tails). For the event of first flipping heads and then tails, the joint probability *p_i_*(*b*, *f*) equals the probability of first flipping heads (0.5) and then tails (0.5), which is 0.5 x 0.5 = 0.25. The individual probabilities *p_i_*(*b*) and *p_i_*(*f*) correspond to the probabilities of getting heads on the first toss (0.5) and tails on the second toss (0.5), respectively. For this state (heads flip and then tails flip), and all the other possible states, the right-hand side of equation (2) will equal 0, because log_2_(1) = 0, and thus the sum of these values also 0.

This example thus yields a mutual information of zero, because the outcome of the first and second coin flip are *independent* of each other.

Now let us assume that for whatever reason, the outcome of the first coin flip somehow influences the outcome of the second coin flip, rendering it more likely to be heads if the first flip yielded heads. In this example, the individual probabilities remain the same, with *p_i_*(*f*) = 0.5 and *p_i_*(*b*) = 0.5, but the joint probabilities differ. Upon completing the calculation in Equation 2, the total mutual information will be greater than 0. The reason is that the two variables are no longer independent. The stronger this statistical dependency is, the greater is the absolute value of the mutual information.

In the context of our experiment, we calculate the mutual information between the identity of different bases at position *i* of a DNA sequence *p_i_*(*b*) and fluorescence scores *p_i_*(*f*). Positions with low mutual information correspond to promoter activity similar to the background, indicating that base identity and fluorescence are independent of each other. In contrast, for positions with high mutual information, some underlying sequence architecture causes fluorescence to be dependent on base identity. Large mutual information indicates that this dependency is strong, for example because position *i* is part of a promoter or a transcription factor binding sites.

### Correcting mutual information calculations for small sample size

Small datasets can skew the mutual information calculation, just as they affect other procedures in statistics. To account for the finite number of mutant daughter sequences that we used to calculate mutual information in Equation 2, we used a previously described correction for finite sample sizes ^31^, which renders the final mutual information we computed equal to equation 3:

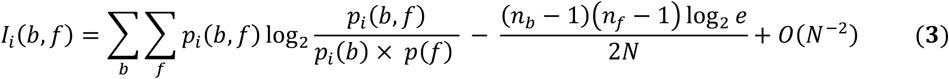

Here *n_b_* is the number of bases (4) and *n_f_* the number of fluorescence bins (4). *N* is the total number of mutant daughter sequences tested. The value O(*N*^−2^) indicates a term that is of the order of *N*^−2^. The correction term is dependent on the degrees of freedom of all possible states (*n_b_* − 1)(*n_f_* − 1) and the size of the library itself. The larger the library, the smaller the correction term.

To visualize mutual information “hotspots,” we additionally smoothened mutual information as a function of position, using a Gaussian filter implemented in the python `scipy` package `ndimage` (parameter alpha=2). We report the mutual information values (smoothened and not smoothened) in **Data S4**.

### Kolmogorov-Smirnov tests

We used a Kolmogorov-Smirnov (KS) test to compare the distribution of promoter signatures along 706 IS3s to a uniform distributions on both the top and bottom DNA strand. It tests the null hypothesis that this distribution is a uniform distribution. For this test, we created a list of promoter signature locations that are normalized for IS3 length, where each data point is the location of an individual promoter signature along one IS3 element, and all data points lie in the interval (0,1). We created individual lists of promoter signatures both for the top and the bottom strand. To generate null uniform distributions, we used the `uniform` function from `scipy.stats` to generate a list of numbers between 0 and 1, with the length of these lists equaling the total number of promoter signatures on the top or bottom strands. We then compared the actual distributions of top or bottom promoter signatures with their respective null distributions using the `kstest` function from `scipy.stats`.

---

**Fig. S1.**
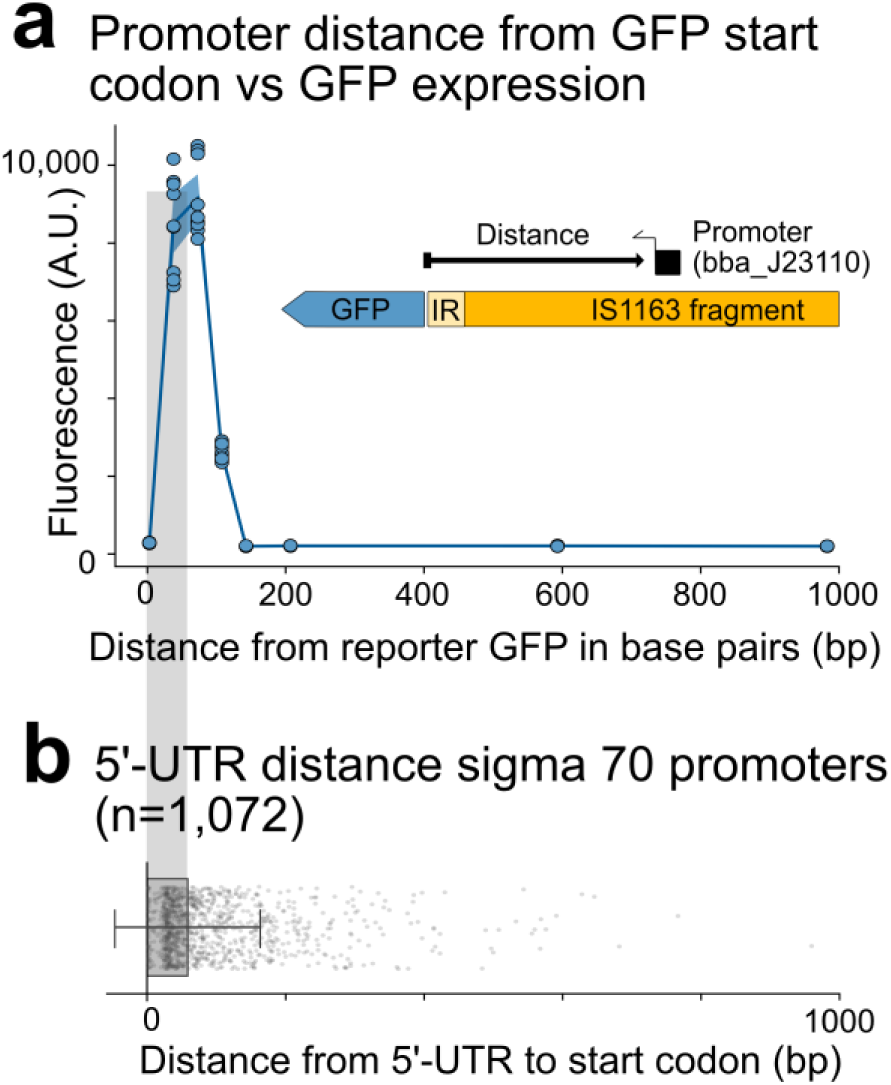
Empirically determining the appropriate construct size for mutation. (**a**) To determine the maximum size of IS fragments allowing the detection of promoter activity within them, we cloned an IS3 fragment (IS1163) downstream of the GFP coding sequence in pMR1. We then placed a moderate-strength constitutive promoter (bba_j23110) at increasing distances from the GFP coding sequence, and measured GFP fluorescence using a plate reader. Circles represent individual measurements, the solid line the mean fluorescence value. The shaded blue area surrounding the mean represents ±1 standard deviation. (**b**) We plot the distance from the 5’ untranslated region (UTR) to the start codon for sigma 70 promoters from the Regulon DB database ^69^. Single points represent individual 5’-UTR lengths. The height of the grey horizontal bar indicates mean 5’-UTR length, and whiskers extend to ±1 standard deviation of the mean.

**Fig. S2.**
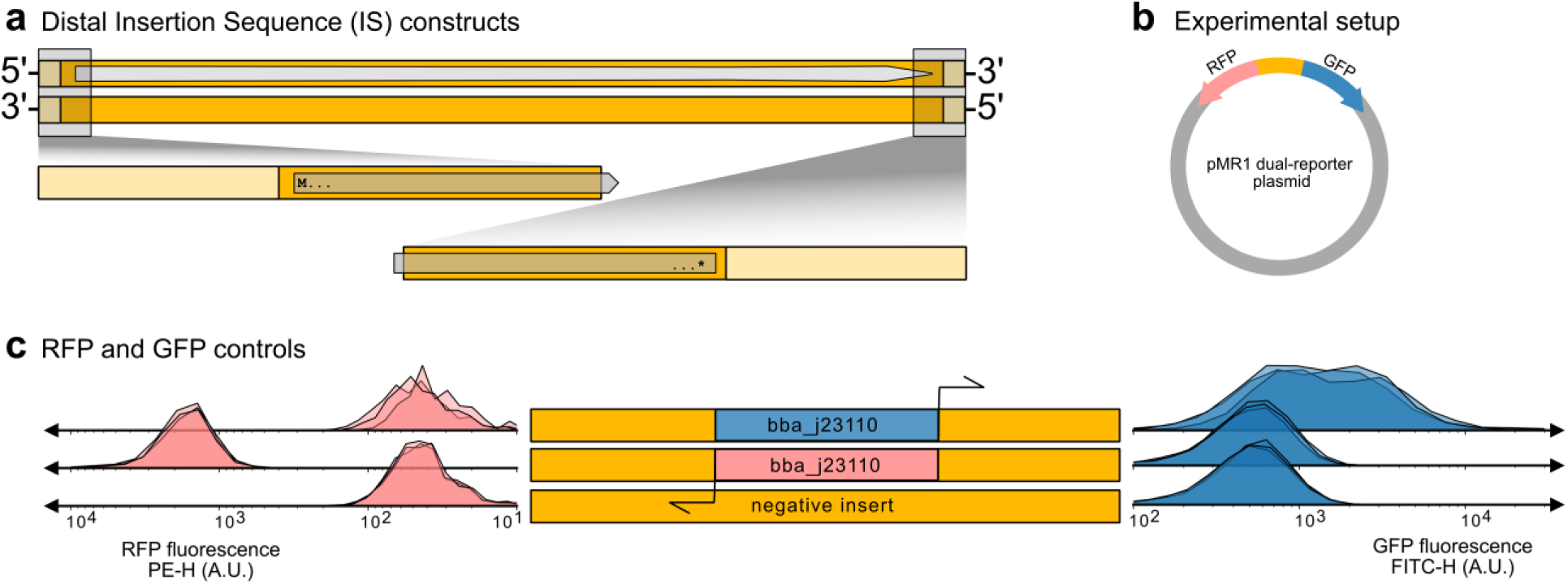
Control constructs for fluorescence readouts. **(a)** We tested the distal ends of IS3s for their ability to drive outward-directed promoter activity. We cloned 120 bps from the ends of each IS3, including the inverted repeat (light yellow) and either the beginning or the end of the transposase coding sequence. **(b)** To test for promoter activity, we cloned the sequences into the pMR1 dual-reporter plasmid between an RFP and GFP coding sequence to simultaneously measure promoter activity in both genetic orientations. The top strand drives GFP expression and the bottom strand RFP expression. **(c)** We compared the promoter activity of the IS3 fragments, as quantified by fluorescence output, to three different controls. For GFP expression, we compared their fluorescence output to a positive control, in which GFP expression is driven by the bba_j23110 promoter (a moderate constitutive promoter) oriented towards the GFP coding sequence. For RFP expression we compared it to fluorescence driven by the bba_j23110 promoter oriented towards the RFP coding sequence. As negative controls, we used the fluorescence readout of pMR1 not encoding an insert, as well as the fluorescence readout of bba_j23110 but oriented in the opposite direction from that in the positive controls.

**Fig. S3.**
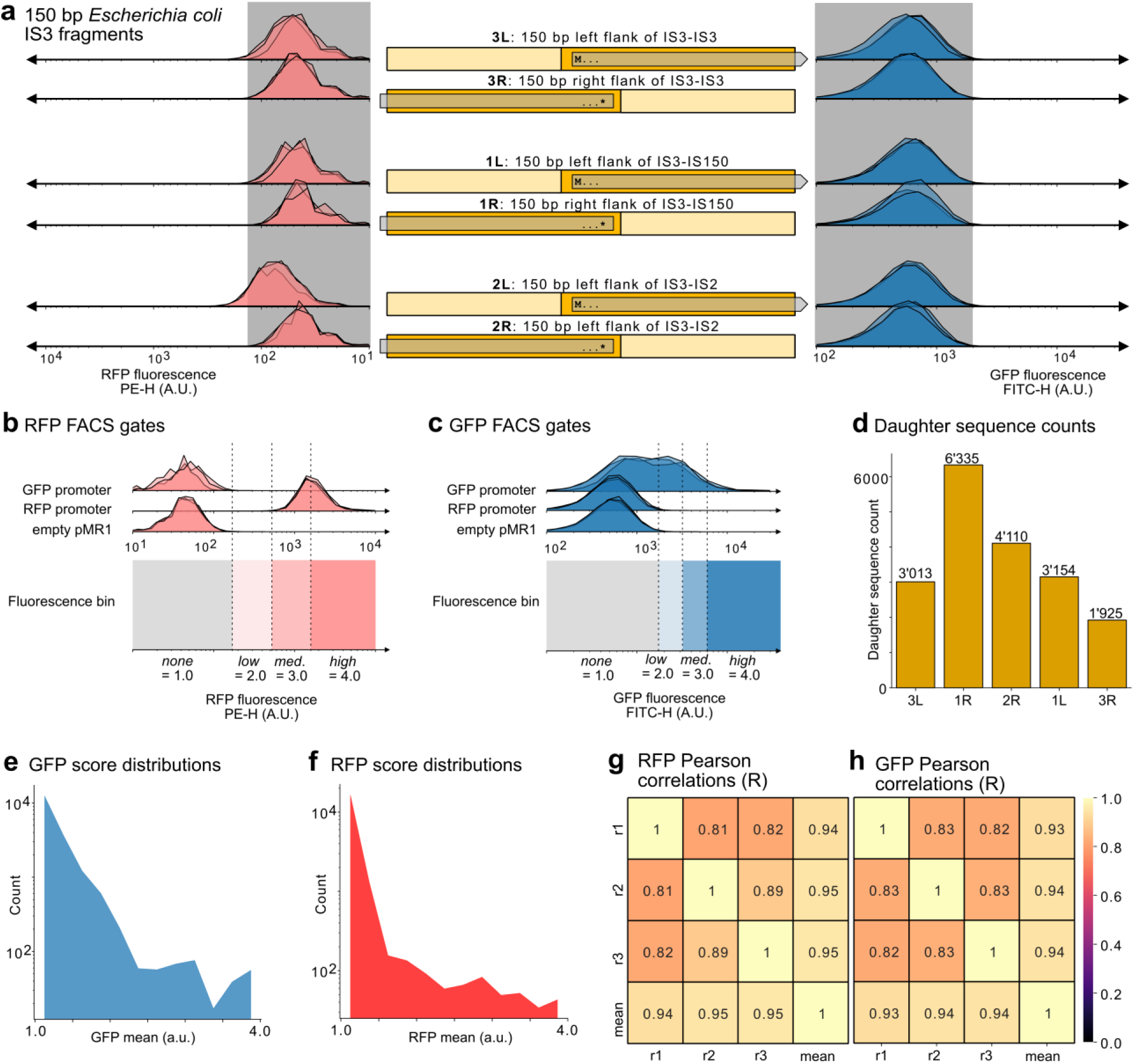
Sort-seq library. **(a)** The IS3 parent sequence fragments used to generate the sort-seq library. For each parent sequence, the middle of the panel represents the parent sequence itself, where the bold name is the shorthand name used throughout this work. “L” denotes the left end of the IS3 and “R” the right end respectively. Light yellow indicates the inverted repeat sequence. The internal gray arrow represents the coding sequence, where “M…” is the start codon and “…*” the stop codon. We measured the promoter activity of each parent sequence in both genetic orientations using a flow cytometer. Left column of histograms: RFP expression, a proxy for the bottom strand’s promoter activity. Right column of histograms: GFP expression, a proxy for the top strand’s promoter activity. The gray box refers to the fluorescence of the negative controls (see also panel **Fig S2c**). We removed sequence 2L from all downstream analysis because it encodes promoter activity on the bottom strand. **(b,c)** We defined fluorescence bin boundaries based on fluorescence measurements from the following three control plasmids. Top: bba j23110 promoter oriented towards GFP. Middle: bba j23110 promoter oriented towards RFP. Bottom: empty pMR1 plasmid. See also **Fig S2c**. See methods for details. See methods for the binning strategy. (**d**) The number of unique mutant “daughter” sequences per “parent” template sequence. **(e,f)** Distribution of fluorescence score (see methods) for GFP **(e)** and RFP **(f).** (**g,h**) Pearson correlation coefficients between the fluorescence scores of the same sequences in different sort-seq replicates (r1, r2, r3), and of the mean fluorescence score (which is the score we used for our analyses) for both RFP (**g**) and GFP (**h**).

**Fig. S4.**
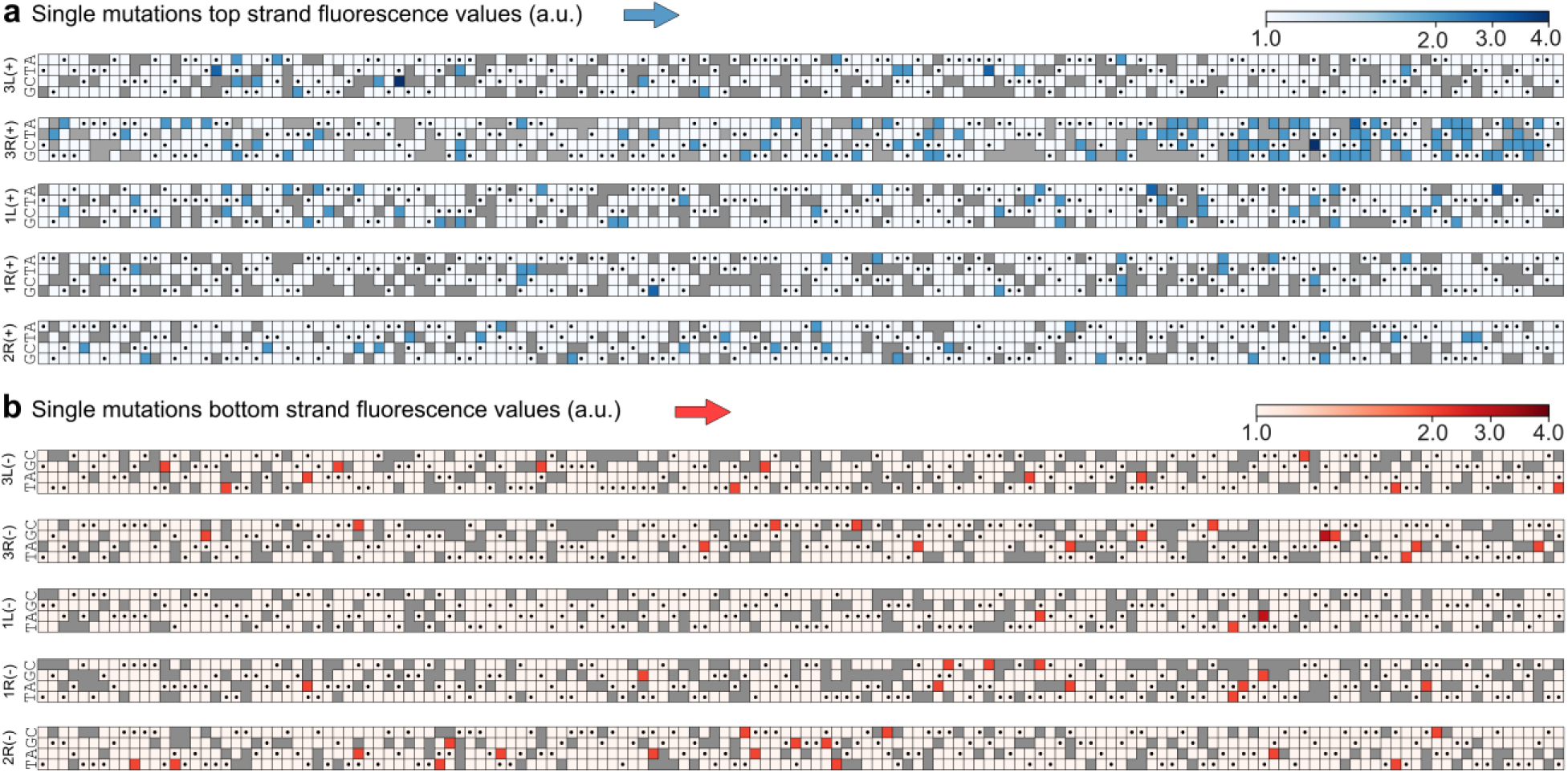
Genotype-phenotype maps for single mutations. (**a,b**) Single mutations observed for each parental IS fragment (rows) and each nucleotide position (columns), together with the new gene expression they drive (blue or red, see color legend). Gray boxes indicate that no mutagenized fragment harbors the indicated nucleotide. Boxes with black circles indicate the wild-type sequence. Sequences are shown from the 5’ to the 3’ end. (**a**) Expression level of top DNA strand (blue, darker: higher expression). (**b**) Expression level of bottom DNA strand (red, darker: higher expression).

**Fig. S5.**
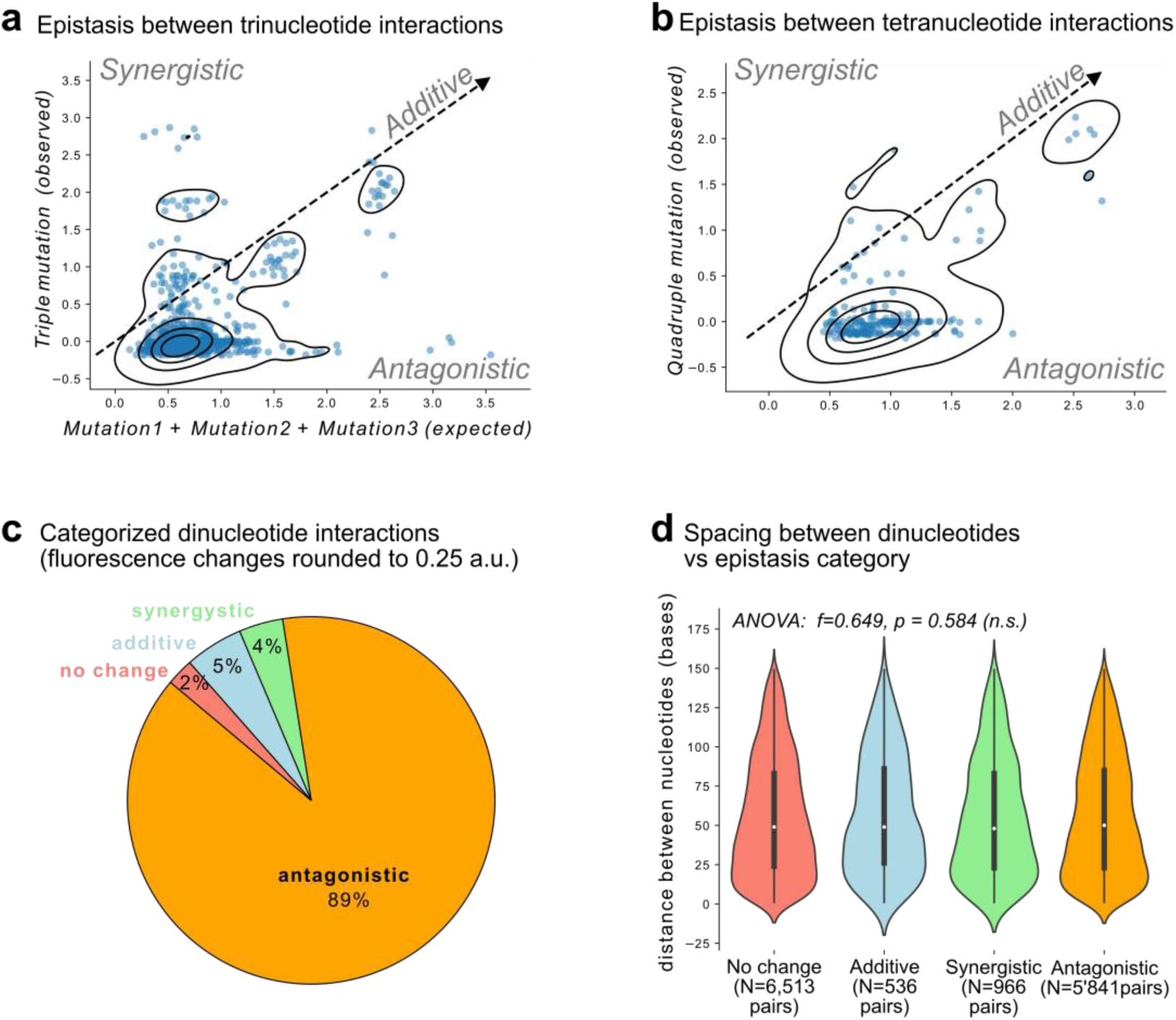
Trinucleotide and tetranucleotide interactions are mostly antagonistic. **(a)** We identified triple mutants in our dataset where we also have mutational data on the expression effects of the constituent individual mutants. We plot the fluorescence changes in arbitrary units (a.u.) for the sum of the individual mutational effects on the x-axis as “expected” values against the fluorescence change of their corresponding triple mutant as an “observed” value. The dotted line indicates equality between expected and observed values. It corresponds to additive interactions. Interactions above (below) the dotted line are synergistic (antagonistic), where the sum of mutational effects is greater (smaller) than expected. Black striations represent kernel density estimates of the scatterplot (seaborn.kdeplot) and illustrate the central tendency of the data, which lies at −0.03 a.u. (observed), and +0.64 a.u. (expected). **(b)** Analogous to (a) but for quadruple mutants. Central tendency is −0.02 a.u. observed, and +0.86 a.u. expected. **(c)** We rounded the fluorescence changes of each dinucleotide interaction to the nearest 0.25 a.u. and classified the interactions as follows. If mut 1 = 0.0 a.u., mut 2 = 0.0 a.u., and mut 1+2 = 0.0 a.u., the interaction is classified as *no change.* If mut1 + mut2 = mut1+2, the interaction is classified as *additive.* If mut1 + mut2 > mut1+2, the interaction is classified as *synergistic.* If mut1 + mut2 < mut1+2, the interaction is classified as *antagonistic.* The percentages of the pie chart correspond to the ratio of each category for the 13,856 dinucleotide interactions. **(d)** We categorized the dinucleotide interactions analogously to (c) but rounded the fluorescence changes to the nearest 0.50 a.u. (see Fig 1i). For each category (*no change, additive, synergistic, antagonistic*), we calculated the distance between the double mutations, and tested the null hypothesis that there is no difference between the means of the distances using an analysis of variance (ANOVA, f=0.649, p=0.584, n.s. = not significant).

**Fig. S6.**
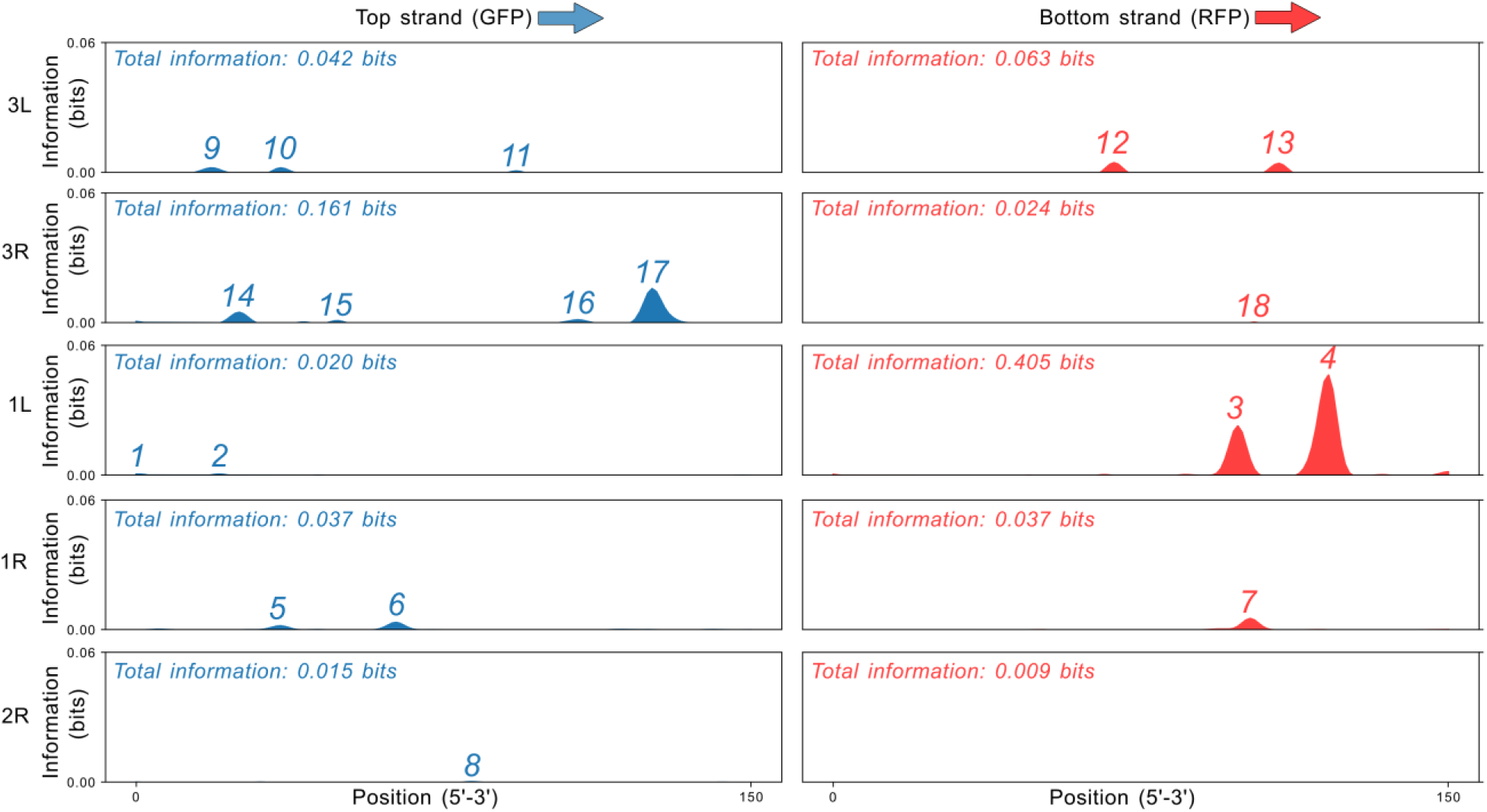
Mutual information for the 10 parent sequences. We calculated the mutual information for each parent sequence (rows) and sequence strand (columns) using Equation 3 (see methods). For each panel the x-axis corresponds to the position (i) in the parent sequence from the 5’ end (base pair 0) to the 3’ end (base pair 150). The y-axes show the amount of mutual information in bits. The heights of the y-axes are fixed to 0.06 bits. Total information equals the sum of mutual information values across all positions.

**Fig. S7.**
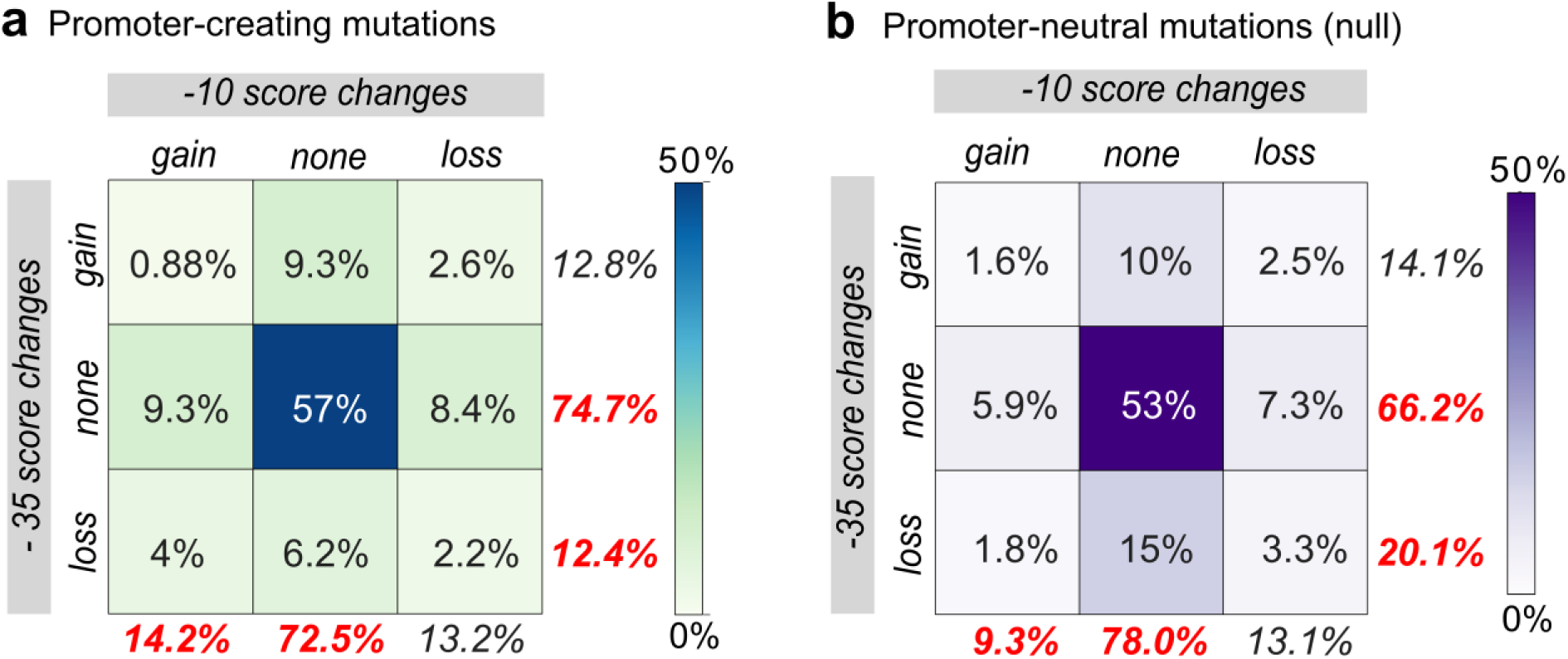
Single mutation biases and contingency tables. (**a,b**) For each point mutation, we calculated how the mutation changed the −10 and −35 position weight matrix (PWM) scores, and classified the changes into either an increase (“gain”), a decrease (“loss”), or no change (“none”) in the PWM score for both the −10 and −35 boxes. We plotted these categories in a contingency matrix. We split the matrix into two groups of changes, those that created weak promoter activity (**a**), and those that did not create promoter activity (promoter-neutral mutations) (**b**). We used a chi-squared test of the null hypothesis that there is no difference between the two contingency tables in (a) and (b). This test rejects that null hypothesis at p=0.014 (4 d.f.). We highlight that −10 scores increase in 14.2% of promoter-creating mutations vs 9.3% in promoter-neutral mutations. We also highlight that −35 scores decrease in 12.4% of cases in promoter-creating mutations, but in 20.1% of promoter-neutral mutations. Remarkably, −10 and −35 scores do not change in promoter-creating mutations 72.5% and 74.7% of the time, respectively, compared to 78.0% and 66.2% of the time in promoter-neutral mutations. Taken together, these numbers suggest that gain or loss of −10 and −35 sites, as indicated by their changing PWM scores, is not the primary path towards weak promoter emergence.

**Fig. S8.**
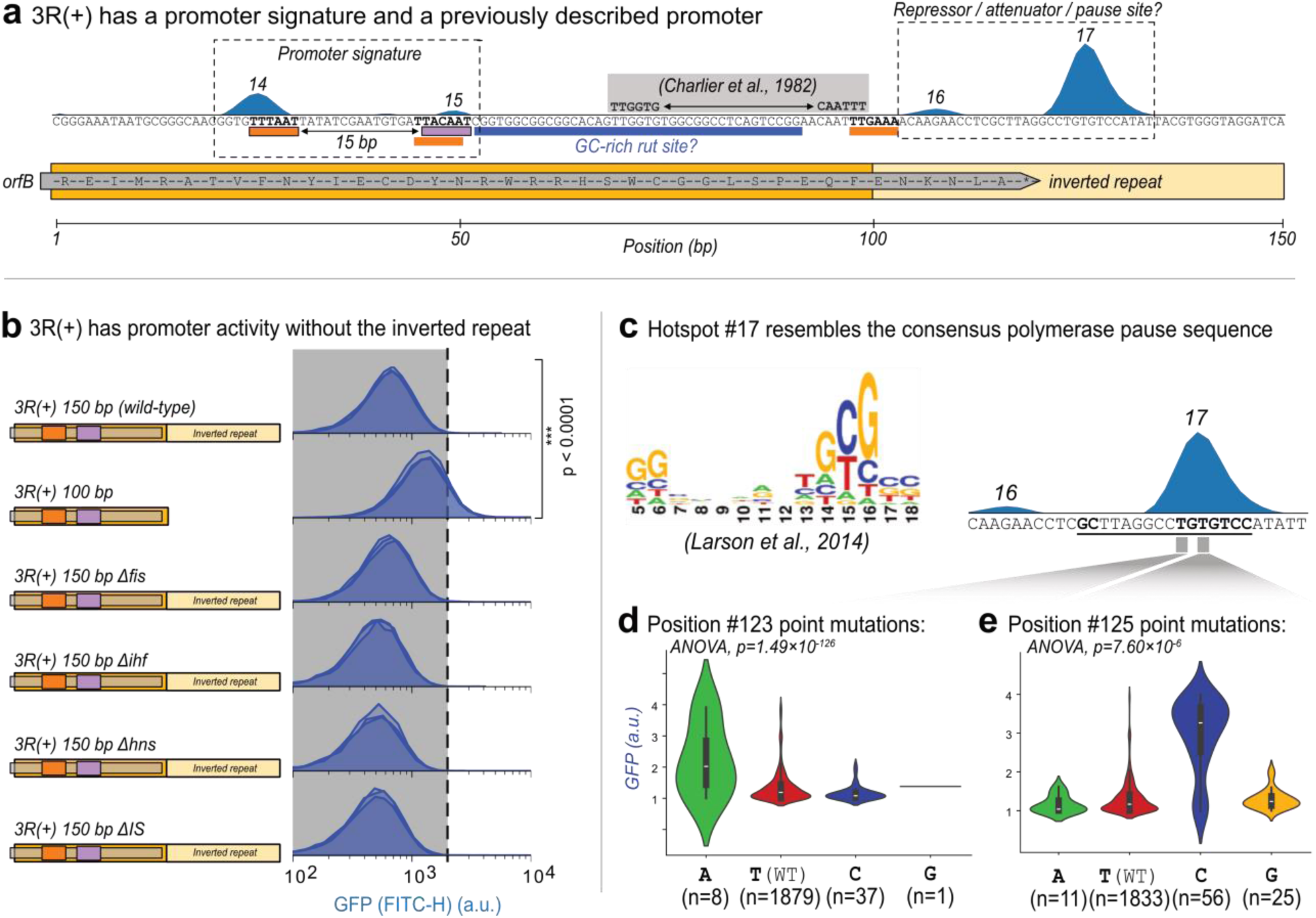
3R(+) may contain a polymerase pause site. **(a)** Parent 3L(+). Top: Mutual information between fluorescence scores and nucleotide identity at every position in the wild-type sequence. The numbers above each peak correspond to their “hotspot” identifiers. Middle: position-weight matrix (PWM) predictions for −10 boxes (magenta rectangles) and −35 boxes (orange rectangles). The gray rectangle corresponds to a previously characterized promoter in ref ^16^. The blue rectangle corresponds to a possible GC-rich Rho utilization site (rut). Bold sequences overlap with either a −10 or −35 box. Bottom: positions along the 3R(+) construct, and its corresponding coding sequence in *orfB.* **(b)** Reporter constructs and their fluorescence readout measured with a flow cytometer. We compared the fluorescence of the wild-type 150 bp 3R(+) parent sequence to the following (from top to bottom): a 100 bp 3R(+) parent sequence without inverted repeats and candidate repressor site (median fluorescence 1’212 a.u. vs 638 a.u., respectively; two-tailed t-test, p=4.3×10-188); the wild-type 3R(+) construct in a Δfis background, a Δihf background, and a Δhns background; as well as the “clean genome” described in ref^59^, which does not contain any insertion sequences (ΔIS). The dashed vertical line and shaded area correspond to the fluorescence values of the wild-type 3R(+) construct in a wild-type (DH5α) background. **(c)** Left: the consensus logo for a polymerase pause sequence identified in ref^61^. Right: DNA sequence and mutual information of 3R(+) as described in (a). The bold underlined region corresponds to the presumed polymerase pause site. The gray regions correspond to regions of interest for the subsequent figure panels. **(d)** GFP fluorescence scores of daughter sequences with point mutations at position #123 of 3R(+). We tested the null hypothesis that the fluorescence distributions are the same for each possible mutation (ANOVA, p=1.49×10-126). **(e)** Analogous to (d) but for point mutations at position #125 (ANOVA, p=7.60×10-6).

**Fig. S9.**
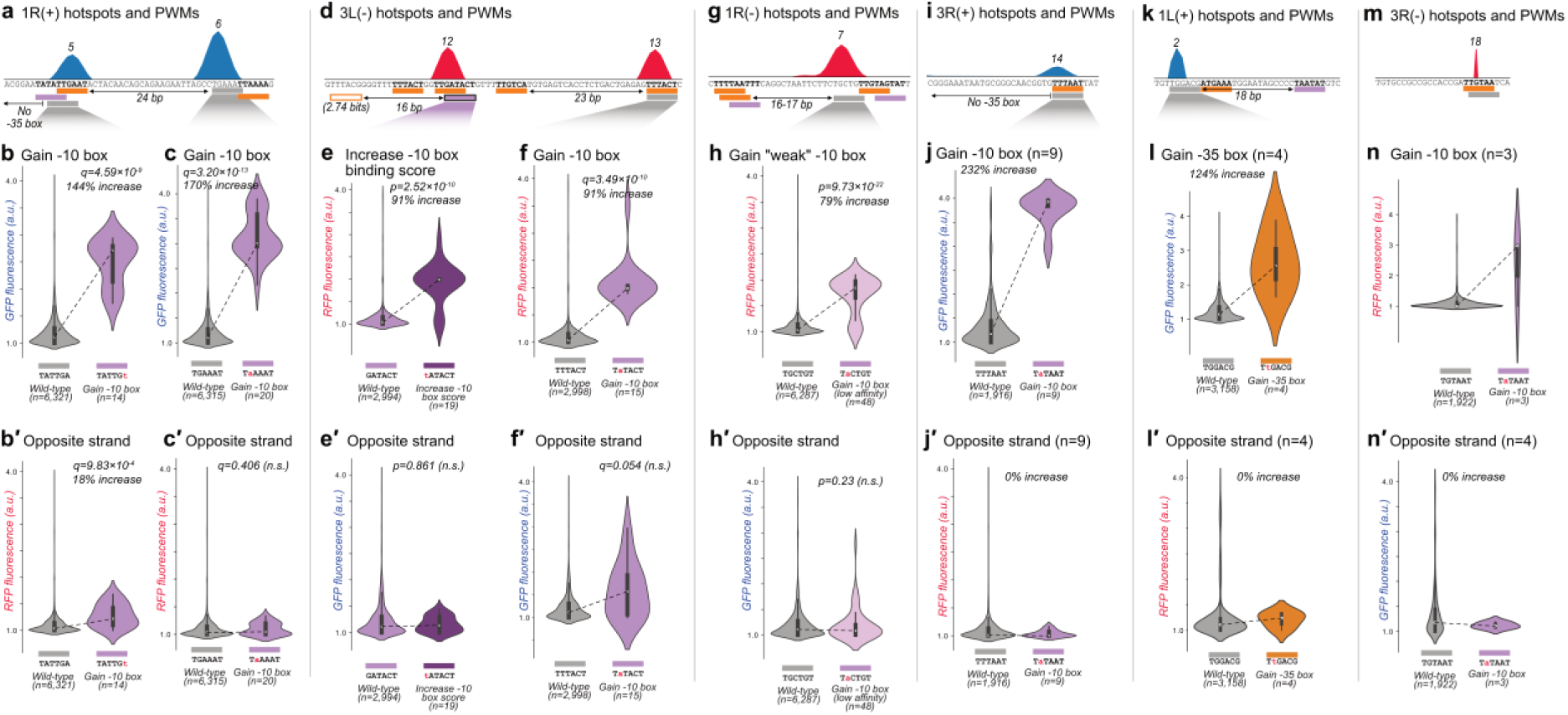
Gaining −10 boxes is associated with increased promoter activity. **(a)** Parent 1R(+). Top: Mutual information between fluorescence scores and nucleotide identity at every position in the wild-type sequence. The numbers above each peak correspond to their hotspot identifiers. Bottom: predictions for −10 boxes (magenta rectangles) and −35 boxes (orange rectangles). Gray rectangles correspond to regions of interest. Bold sequences overlap with either a −10 or −35 box. Arrows indicate the distance between the regions of interest to the nearest −10 or −35 box. **(b)** GFP fluorescence scores for daughters without vs with a −10 box in the region of interest shown in panel (a). The most frequent genotype is written below for each category. We test the null hypothesis that gaining a −10 box in this region of interest does not increase fluorescence (two-tailed Mann-Whitney U [MWU] test), and correct the p-values using a Benjamini-Hochberg correction as corresponding q-values (see methods, two-tailed MWU test, q=4.59×10-9). The area of each violin plot corresponds to the kernel density estimate of each distribution. Within each violin plot is a boxplot where the white circle corresponds to the median, the box the interquartile range (IQR), and the whiskers 1.5×IQR. **(b′)** Like (b) but for RFP fluorescence scores on the opposite strand (two-tailed MWU test, q=9.83×10-4). **(c)** Like (b) but for the other region of interest shown in (a) (two-tailed MWU test, q=3.20×10-13). **(c′)** Like (c) but for the RFP fluorescence scores on the opposite strand (two-tailed MWU test, q=0.406, n.s.=not significant). **(d)** Like (a) but for parent 3L(−). **(e)** RFP fluorescence scores for daughters with a wild-type −10 box vs a −10 box with a PWM score greater than the wild-type box. We test the null hypothesis that increasing the PWM score of the −10 box in this region does not increase fluorescence (two-tailed MWU test, p=2.52×10-10). This was a post-hoc test and we therefore do not Benjamini-Hochberg correct the p-value (see methods). **(e′)** Like (e) but for the GFP fluorescence scores on the opposite strand (two-tailed MWU test, p=0.861, n.s.=not significant). **(f)** Like (b) but for the other region of interest shown in (d) (two-tailed MWU test, q=3.49×10-10). **(f′)** Like (f) but for the GFP fluorescence scores on the opposite strand (two-tailed MWU test, q=0.054, n.s.=not significant). **(g)** Like (a) but for Parent 1R(−). **(h)** RFP fluorescence scores for daughters without a −10 box in the region of interest vs daughters with a “low-affinity” −10 box (PWM threshold = 1.99 bits instead of 3.98 bits). We test the null hypothesis that gaining this low affinity −10 box does not increase fluorescence (two-tailed MWU test, p=9.73×10-22). This was a post-hoc test and we therefore do not correct the p-value using Benjamini-Hochberg (see methods). **(h′)** Like (h) but for the GFP fluorescence scores on the opposite strand (two-tailed MWU test, p=0.23, n.s.=not significant). **(i-n′)** For these hotspots, one of the two groups being compared has fewer than 10 samples (n<10), and we cannot reliably report p or q-values from two-tailed MWU tests. **(i)** Analogous to (a) but for Parent 3R(+). **(j)** Analogous to (b) but for the region of interest shown in (i). **(j′)** Like (j) but for the RFP fluorescence scores on the opposite strand. **(k)** Analogous to (a) but for Parent 1L(+). **(l)** Like (b) but for the region of interest shown in (k) and for a −35 box PWM instead of a −10 box PWM. **(l′)** Like (l) but for the RFP fluorescence scores on the opposite strand. **(m)** Like (a) but for Parent 3R(−). **(n)** Like (b) but for the region of interest shown in (m). **(n′)** Analogous to the mutations and descriptions in (n), but for GFP fluorescence scores on the opposite strand.

**Fig. S10.**
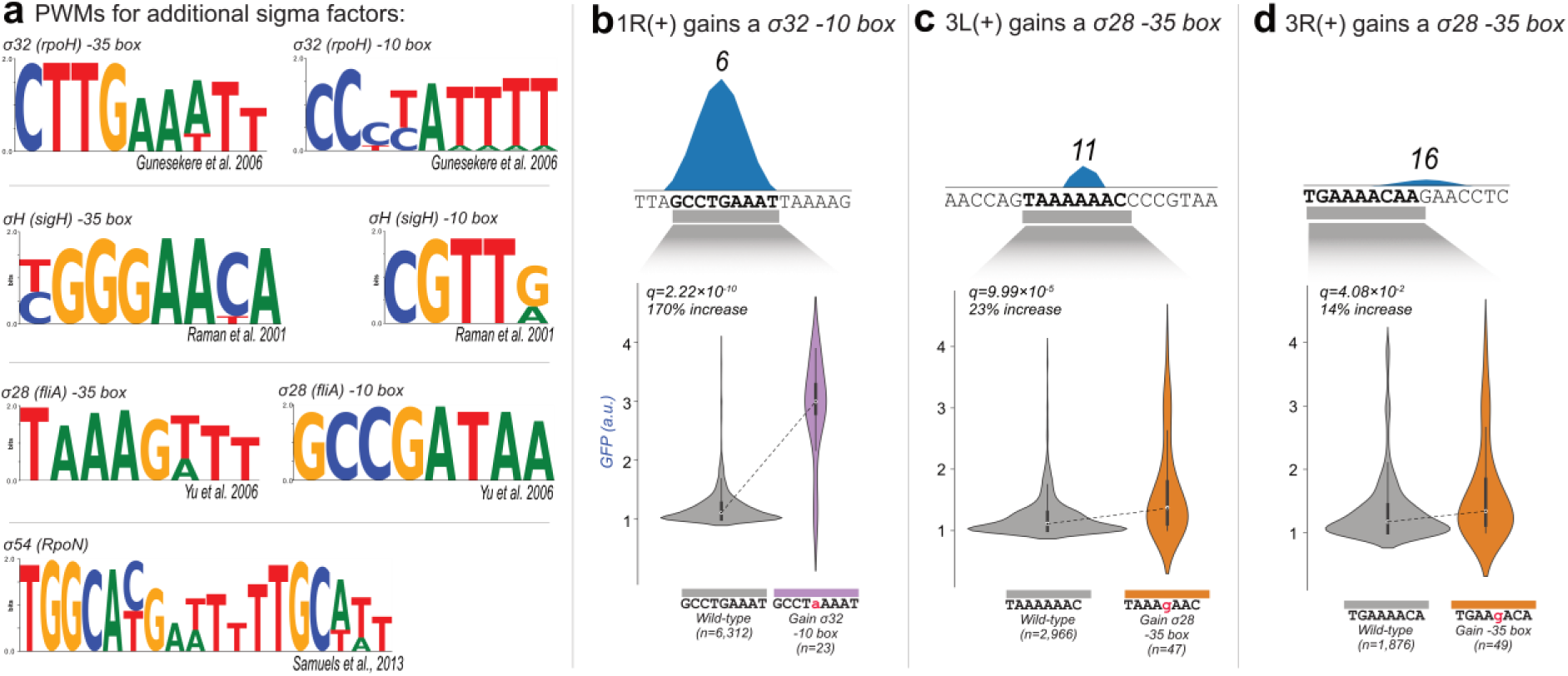
Gaining motifs for other sigma factors. **(a)** We acquired position weight matrices (PWMs) for additional sigma (σ) factors. Specifically, we acquired the σ32 −35 and −10 box PWMs from ref^72^, σH −35 and −10 box PWMs from ref^73^, σ28 −35 and −10 boxes from ref^74^, and σ54 from ref^75^. Logos were drawn using Logomaker^76^. See methods. **(b)** Parent 1R(+). Top: Mutual information between fluorescence scores and nucleotide identity at every position in the wild-type sequence. The numbers above each peak correspond to their hotspot identifiers. Gray rectangles correspond to regions of interest. Bottom: GFP fluorescence scores for daughters without vs with a σ32 −10 box in the region of interest. The most frequent genotype is written below for each category, with the difference between the wild-type sequence written in red, lowercase font. We test the null hypothesis that gaining a σ32 −10 box in this region of interest does not increase fluorescence (two-tailed Mann-Whitney U [MWU] test), and correct the p-values using a Benjamini-Hochberg correction as corresponding q-values (see methods, two-tailed MWU test, q=2.22×10-10). The area of each violin plot corresponds to the kernel density estimate of each distribution. Within each violin plot is a boxplot where the white circle corresponds to the median, the box to the interquartile range (IQR), and the whiskers to 1.5×IQR. **(c)** Like (b) but for gaining a σ28 −35 box in 3L(+) (two-tailed MWU test, q=9.99×10^−5^). **(d)** Like (b) but for gaining a σ28 −35 box in 3R(+) (two-tailed MWU test, q=4.08×10-2).

**Fig. S11.**
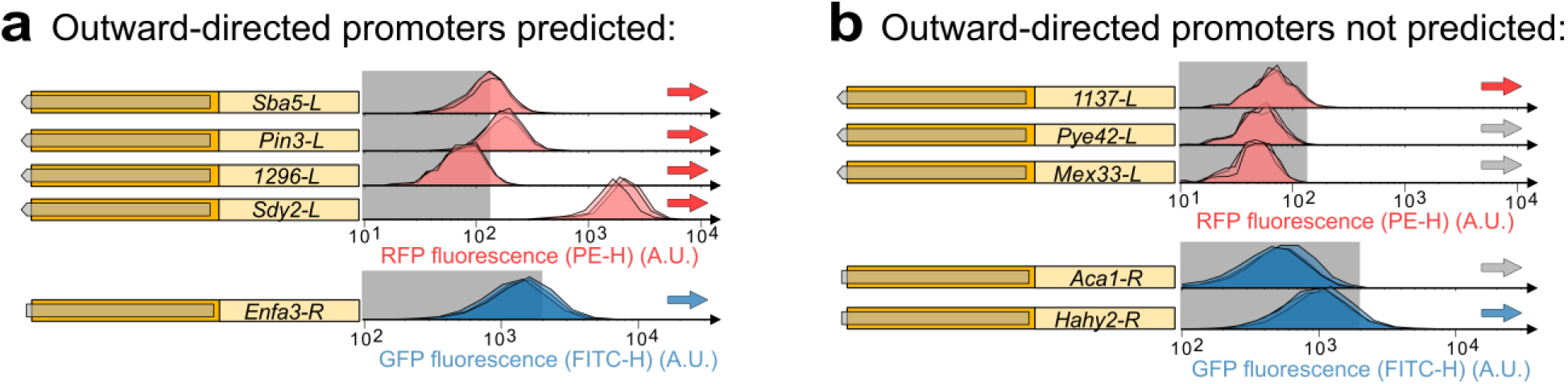
testing promoter signatures for promoter activity. **(a,b)** Flow cytometry plots depicting the distribution of fluorescence as arbitrary units (a.u.) for genetic constructs cloned into a reporter plasmid (pMR1). Depending on the orientation, the fluorescence readout is either for RFP (red histograms, bottom strand) or GFP (blue histograms, top strand). Shaded regions correspond to readouts produced by negative controls (see methods). Blue and red arrows indicate high fluorescence, and gray arrows indicate fluorescence indistinguishable from controls. **(a)** Left: fluorescence readouts for five randomly selected ends of IS3 sequences predicted to have outward-directed promoter activity, and right: **(b)** for five randomly selected predicted not to have outward promoter signatures.

**Figure S12.**
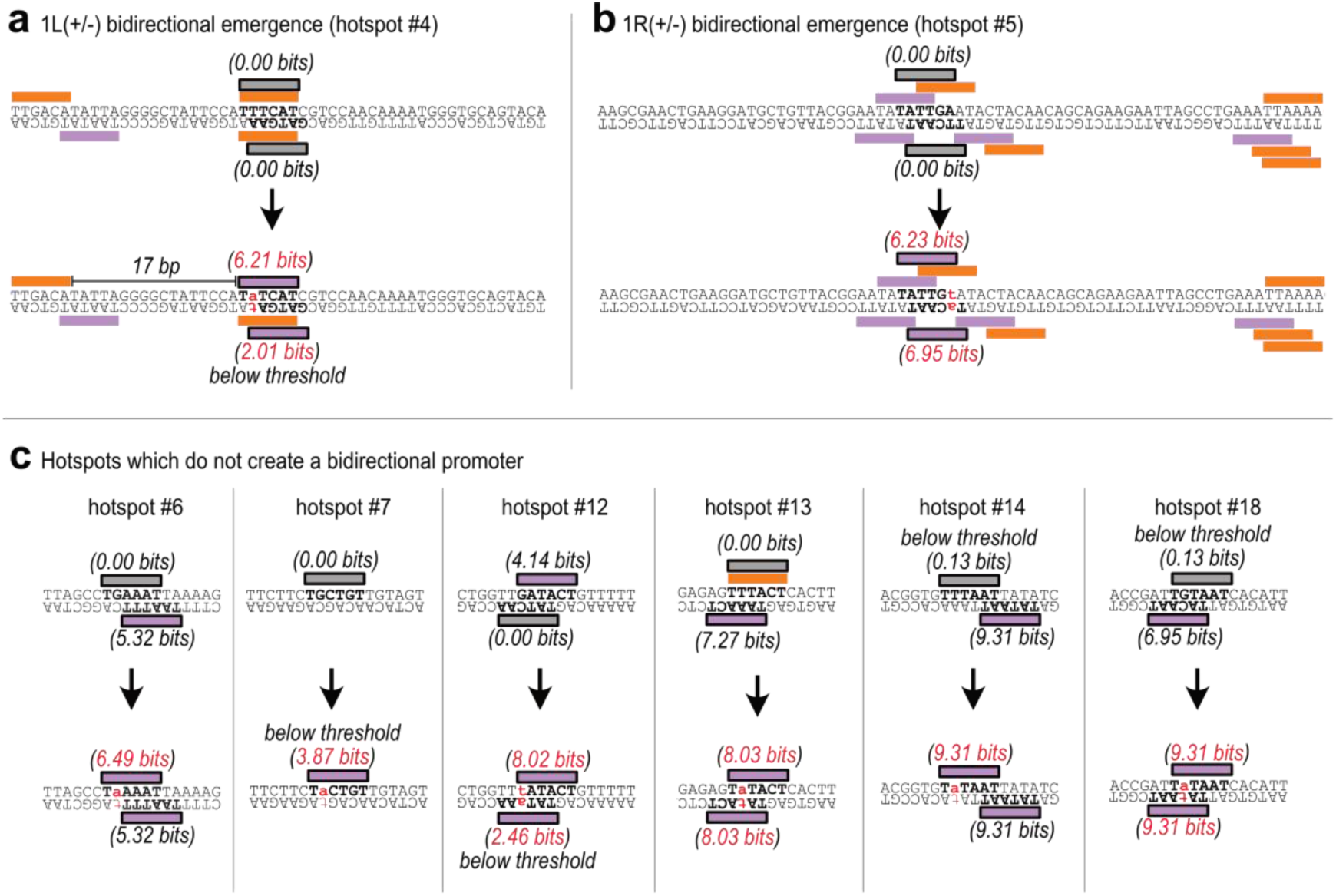
Gaining symmetrical −10 boxes is not sufficient to create a bidirectional promoter. **(a)** Double-stranded parent DNA sequence for parent 1L(+/−). Orange rectangles correspond to −35 boxes and magenta rectangles to −10 boxes. Gray boxes correspond to the region of interest. Numbers in Information theoretical units (bits) are position-weight matrix (PWM) scores for the −10 boxes. Top: the wild-type parent sequence without bidirectional promoter activity. Bottom: the point mutation that creates a bidirectional promoter is shown in red, lowercase font. **(b)** Analogous to (a) but for parent 1R(+/−). **(c)** Analogous to (a) but for hotspots that do not create bidirectional promoters when mutations create a −10 box on the top strand.

**Table S1.** Sorting events and replicates. All information on the number of events sorted for each replicate and day during fluorescence activated cell soring (FACS)

|  | GFP |  |  |  | RFP |  |  |  |
| --- | --- | --- | --- | --- | --- | --- | --- | --- |
| Bin | 1 | 2 | 3 | 4 | 1 | 2 | 3 | 4 |
| Sort day 1 | 3.6e+6<br>x1<br>replicate | 4.5e+5<br>x1<br>replicate | 3.0e+5<br>x1<br>replicate | 13'045<br>x1<br>replicate | 3.6e+6<br>x1<br>replicate | 4.5e+5<br>x1<br>replicate | 33'681<br>x1<br>replicate | 10,004<br>x1<br>replicate |
| Sort day 2 | 3.6e+6<br>x3<br>replicates | 4.5e+5<br>x3<br>replicates | 3.0e+5<br>x3<br>replicates | 1.5e+5<br>x3<br>replicates | 3.6e+6<br>x3<br>replicates | 4.5e+5<br>x3<br>replicates | 1.0e+5<br>x3<br>replicates | 1.0e+5<br>x3<br>replicates |

**Table S2.**
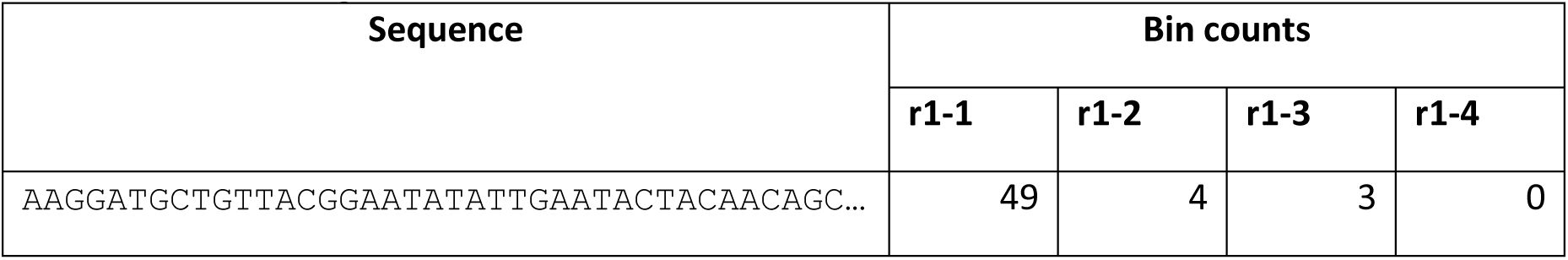
Calculating fluorescence scores. An example dataset to illustrate how we calculated final fluorescence scores. See Methods subsection: processing sequencing results for equation (1).

